# Evolutionary origins of the SARS-CoV-2 sarbecovirus lineage responsible for the COVID-19 pandemic

**DOI:** 10.1101/2020.03.30.015008

**Authors:** Maciej F Boni, Philippe Lemey, Xiaowei Jiang, Tommy Tsan-Yuk Lam, Blair Perry, Todd Castoe, Andrew Rambaut, David L Robertson

**Affiliations:** Center for Infectious Disease Dynamics, Department of Biology, Pennsylvania State University, University Park, PA, USA; Department of Microbiology, Immunology and Transplantation, KU Leuven, Leuven, Belgium; Xi’an Jiaotong-Liverpool University (XJTLU), Suzhou, Jiangsu, China; State Key Laboratory of Emerging Infectious Diseases, School of Public Health, The University of Hong Kong, Hong Kong SAR, China; Department of Biology, University of Texas Arlington, Arlington, TX, USA; Institute of Evolutionary Biology, University of Edinburgh, Edinburgh, UK; MRC-University of Glasgow Centre for Virus Research (CVR), Glasgow, UK

## Abstract

There are outstanding evolutionary questions on the recent emergence of coronavirus SARS-CoV-2/hCoV-19 in Hubei province that caused the COVID-19 pandemic, including (1) the relationship of the new virus to the SARS-related coronaviruses, (2) the role of bats as a reservoir species, (3) the potential role of other mammals in the emergence event, and (4) the role of recombination in viral emergence. Here, we address these questions and find that the sarbecoviruses – the viral subgenus responsible for the emergence of SARS-CoV and SARS-CoV-2 – exhibit frequent recombination, but the SARS-CoV-2 lineage itself is not a recombinant of any viruses detected to date. In order to employ phylogenetic methods to date the divergence events between SARS-CoV-2 and the bat sarbecovirus reservoir, recombinant regions of a 68-genome sarbecovirus alignment were removed with three independent methods. Bayesian evolutionary rate and divergence date estimates were consistent for all three recombination-free alignments and robust to two different prior specifications based on HCoV-OC43 and MERS-CoV evolutionary rates. Divergence dates between SARS-CoV-2 and the bat sarbecovirus reservoir were estimated as 1948 (95% HPD: 1879-1999), 1969 (95% HPD: 1930-2000), and 1982 (95% HPD: 1948-2009). Despite intensified characterization of sarbecoviruses since SARS, the lineage giving rise to SARS-CoV-2 has been circulating unnoticed for decades in bats and been transmitted to other hosts such as pangolins. The occurrence of a third significant coronavirus emergence in 17 years together with the high prevalence and virus diversity in bats implies that these viruses are likely to cross species boundaries again.

**In Brief:** The Betacoronavirus SARS-CoV-2 is a member of the sarbecovirus subgenus which shows frequent recombination in its evolutionary history. We characterize the extent of this genetic exchange and identify non-recombining regions of the sarbecovirus genome using three independent methods to remove the effects of recombination. Using these non-recombining genome regions and prior information on coronavirus evolutionary rates, we obtain estimates from three approaches that the most likely divergence date of SARS-CoV-2 from its most closely related available bat sequences ranges from 1948 to 1982.

**Key Points:**

- RaTG13 is the closest available bat virus to SARS-CoV-2; a sub-lineage of these bat viruses is able to infect humans. Two sister lineages of the RaTG13/SARS-CoV-2 lineage infect Malayan pangolins.
- The sarbecoviruses show a pattern of deep recombination events, indicating that there are high levels of co-infection in horseshoe bats and that the viral pool can generate novel allele combinations and substantial genetic diversity; the sarbecoviruses are efficient ‘explorers’ of phenotype space.
- The SARS-CoV-2 lineage is not a recent recombinant, at least not involving any of the bat or pangolin viruses sampled to date.
- Non-recombinant regions of the sarbecoviruses can be identified, allowing for phylogenetic inference and dating to be performed. We constructed three such regions using different methods.
- We estimate that RaTG13 and SARS-CoV-2 diverged 40 to 70 years ago. There is a diverse unsampled reservoir of generalist viruses established in horseshoe bats.
- While an intermediate host responsible for the zoonotic event cannot be ruled out, the relevant evolution for spillover to humans very likely occurred in horseshoe bats.

## Introduction

In December 2019, a cluster of pneumonia cases epidemiologically linked to an open-air wet market in city of Wuhan (Hubei Province), China (Li et al., 2020a; Zhou et al., 2020b) led local health officials to issue an epidemiological alert to the Chinese Center for Disease Control and Prevention (China CDC) and the World Health Organization’s (WHO) China Country Office. In early January, the etiological agent of the pneumonia cases was found to be a coronavirus (World Health Organization, 2020a), subsequently named SARS-CoV-2 by an International Committee on Taxonomy of Viruses (ICTV) Study Group (Gorbalenya et al., 2020), (also named hCoV-19 by (Wu et al., 2020b)). The first available sequence data (Wu et al., 2020a) placed this novel human pathogen in the *Sarbecovirus* subgenus of *Coronaviridae* (Lu et al., 2020), the same sub-genus as the SARS virus that caused a global outbreak of nearly 8000 cases in 2002-2003. By mid-January, the virus was spreading widely inside Hubei province and by early March SARS-CoV-2 reached pandemic status (World Health Organization, 2020b).

In outbreaks of zoonotic pathogens, identification of the infection source is crucial, as this may allow health authorities to separate human populations from the wild-life or domestic-animal reservoirs posing the zoonotic risk (Stegeman et al., 2004; Yu et al., 2013). If controlling an outbreak in its early stages is not possible – as was the case for the COVID-19 epidemic in Hubei province – identification of origins and point sources is nevertheless important for containment purposes in other provinces and prevention of future outbreaks. When the first genome sequence of SARS-CoV-2, Wuhan-Hu-1, was released on January 10 2020 on Virological.org by a consortium led by Yong-Zhen Zhang (Wu et al., 2020a) it enabled the immediate analyses of its ancestry. Across a large region of its genome, corresponding approximately to ORF1b, it did not cluster with any of the known bat coronaviruses indicating that recombination likely played a role in the evolutionary history of these viruses (Lu et al., 2020; Wu et al. 2020b). Subsequently, a bat sarbecovirus – RaTG13 sampled from a *Rhinolophus affinis* horseshoe bat in 2013 in Yunnan province – was reported that clusters with SARS-CoV-2 in almost all genomic regions with approximately 96% genome sequence identity (Zhou et al., 2020b). Zhou et al. (2020b) concluded from the genetic proximity of SARS-CoV-2 to a bat virus that a bat origin for the current COVID-19 outbreak is probable. Recent evidence has also identified pangolins as potential intermediate species for SARS-CoV-2 emergence or as a potential reservoir species themselves (Lam et al., 2020; Xiao et al., 2020).

Unlike other viruses that have emerged in the past two decades, coronaviruses are highly recombinogenic (Forni et al., 2017; Hon et al., 2008; Lam et al., 2018). Influenza viruses reassort (Webster et al., 1992) but they do not undergo homologous recombination within RNA segments (Boni et al., 2008, 2010), meaning that ‘origins’ questions for influenza outbreaks can always be reduced to origins questions for each of influenza’s eight RNA segments. For coronaviruses however, recombination means that small genomic sub-regions can have independent origins, identifiable if sufficient sampling has been done in the animal reservoirs that support the endemic circulation, coinfection, and recombination that appear to be common. Here, we analyse the evolutionary history of SARS-CoV-2, using available genomic data on sarbecoviruses. We demonstrate that the sarbecoviruses circulating in horseshoe bats have complex recombination histories permitting generation of novel viral variants, as reported by others (He et al., 2014; Hon et al., 2008; Hu et al., 2017; Li et al., 2020b; Lin et al., 2017; Wang et al., 2017; Yuan et al., 2010; Zhou et al., 2020a). Interestingly, despite the lineage leading to SARS-CoV-2 acquiring residues in its Spike (S) protein’s receptor-binding domain (RBD) permitting use of human ACE2 (Wan et al., 2020) and being closer to the pangolin virus than RaTG13 in this region (Lam et al., 2020) – a signal that suggests recombination – the divergence patterns in the S protein show no evidence of recombination between SARS-CoV-2 and known sarbecoviruses. Our results indicate the presence of a single lineage circulating in bats with properties that allowed it to infect human cells, as previously described for the bat sarbecoviruses related to the first SARS-CoV lineage (Ge et al., 2013; Menachery et al., 2015, 2016).

To gauge the length of time this lineage has circulated in bats we estimate the time to most recent common ancestor (tMRCA) of SARS-CoV-2 and RaTG13. We use three bioinformatic approaches to remove the effects of recombination, and we combine these approaches to identify putative non-recombinant regions that can be used for reliable phylogenetic reconstruction and dating. Collectively our analyses point to bats being the primary reservoir for the SARS-CoV-2 lineage. While it is possible that pangolins may have acted as an intermediate species facilitating transmission to humans, the evidence is consistent with the virus having evolved in bats resulting in bat sarbecoviruses that can replicate in the upper respiratory tract of both humans and pangolins (Zhang and Holmes, 2020; Zhou et al., 2020a).

## Results

### Recombination analysis and identification of breakpoint-free genome regions

Of the 68 sequences in the aligned sarbecovirus sequence set, 67 show evidence of mosaicism, indicating involvement in homologous recombination either directly with identifiable parentals or in their deeper shared evolutionary history, i.e., due to shared ancestral recombination events (all Dunn-Sidak corrected *p* < 4 × 10^−4^, 3SEQ (Lam et al., 2018)). This is evidence for numerous recombination events occurring in the evolutionary history of the sarbecoviruses (Li et al., 2020b); however, identifying all past events and their temporal order (Eden et al., 2013) is challenging. Figure 1 (top) shows the distribution of all identified breakpoints (using 3SEQ’s exhaustive triplet search) by the number of candidate recombinant sequences supporting them. The histogram allows for the identification of non-recombining regions by revealing regions with no breakpoints. Sorting these breakpoint-free regions (BFRs) by length results in two segments longer than 5kb: an ORF1a sub-region spanning nucleotides 3625-9150 and the first half of ORF1b spanning nucleotides 13291-19628. Eight other BFRs longer than 500nt were identified. Of the nine breakpoints (defining these ten BFRs), four showed phylogenetic incongruence (PI) signals with bootstrap support higher than 80%, adopting previously published criteria on using a combination of mosaic and PI-signals to show evidence of past recombination events (Boni et al., 2010). All four of these breakpoints were also identified with the tree-based recombination detection method GARD (Kosakovsky Pond et al., 2006).

**Figure 1.**
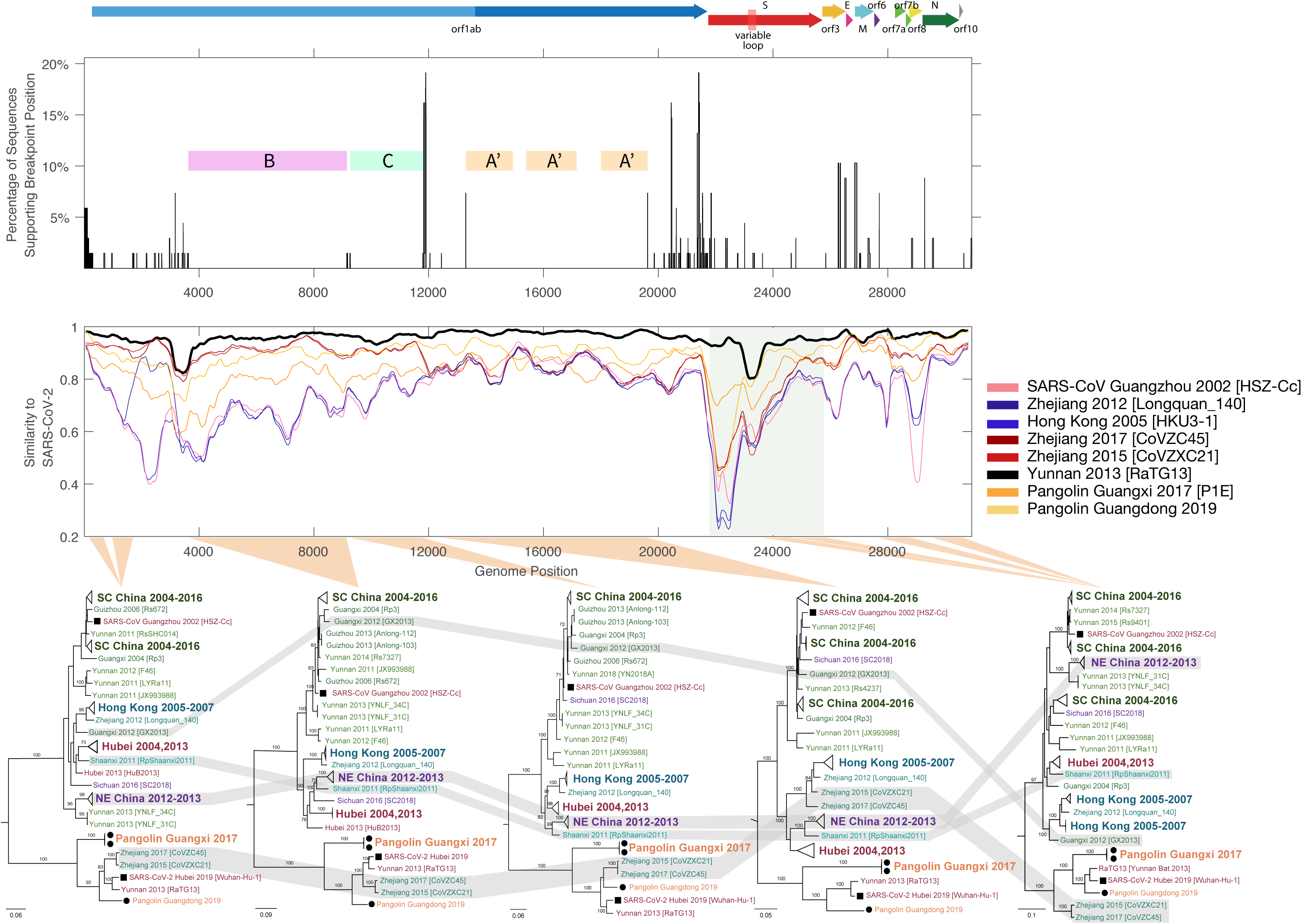
<u>Top:</u> Breakpoints identified by 3SEQ shown by the percentage of sequences (out of 68) that support a particular breakpoint position. Note that breakpoints can be shared between sequences if they are descendants of the same recombination events. The pink, green, and orange bars show breakpoint-free regions (BFRs), with Region **A** (nt 13291-19628) showing two trimmed segments to yield Region **A’** (nt 13291-14932, 15405-17162, 18009-19628). Region **B** spans nucleotides 3625-9150, and region **C** spans 9261-11795. Concatenated region **A’BC** is non-recombining region 1 (**NRR1**). Open reading frames are shown above the breakpoint plot, with the variable loop region indicated in the S protein. <u>Center:</u> Similarity plot between SARS-CoV-2 and several selected sequences including RaTG13 (black), SARS 2002-2003 (pink), and two pangolin sequences (orange). Shaded region corresponds to the S protein. <u>Bottom:</u> Maximum likelihood phylogenetic trees rooted on a 2007 virus sampled in Kenya (BtKy72; root truncated from images), shown for five breakpoint-free regions of the sarbecovirus alignment. Nucleotide positions for phylogenetic inference are 147-695, 962-1686 (first tree), 3625-9150 (second tree, also BFR **B**), 9261-11795 (third tree, also BFR **C**), 12443-19638 (fourth tree), and 23631-24633, 24795-25847, 27702-28843, 29574-30650 (fifth tree). Relevant bootstrap values are shown on branches, and gray-shaded regions show sequences that exhibit phylogenetic incongruence along the genome. SC China corresponds to provinces in southcentral China, specifically Yunnan, Guizhou, and Guangxi provinces. NE China corresponds to provinces in northeast China: Jilin, Shaanxi, Shanxi, Hebei, and Henan provinces.

The extent of the sarbecoviruses’ recombination history can be demonstrated by five phylogenetic trees inferred from BFRs or concatenated adjacent BFRs (Figure 1, bottom). BFRs were concatenated if no phylogenetic incongruence signal could be identified between them. When viewing the last 7kb of the genome a clade of viruses from northeast (NE) China appears to cluster with sequences from south-central (SC) Chinese provinces, but when inspecting trees from different parts of ORF1ab the NE China clade is phylogenetically separated from the SC China clade. Individual sequences such as RpShaanxi2011, Guangxi GX2013, and two sequences from Zhejiang province (CoVZXC21/CoVZC45), as previously shown (Li et al., 2020b; Zhou et al., 2020a), show strong phylogenetic recombination signals, as they fall on different evolutionary lineages (with bootstrap support > 80%) depending what region of the genome is being examined.

Despite this high frequency of recombination among bat viruses, the ‘block-like’ nature of the genome permits retrieval of a clean sub-alignment for phylogenetic analysis. Conservatively, we can combine three >2kb BFRs identified above into a putative non-recombining region 1 (**NRR1**), after removing five sequences that appear to be recombinants and two small sub-regions of the longer region (nucleotides 13291-19628). Alternatively, combining 3SEQ-inferred breakpoints with GARD-inferred breakpoints and the necessity of PI-signals for inferring recombination, we can use the 9.9kb region spanning nucleotides 11885-21753 (**NRR2**) as a putative non-recombining region; this approach is conservative in identifying breakpoints but not conservative in identifying non-recombining regions. Using a third consensus-based approach for identifying recombinant regions in individual sequences – with six different recombination detection methods in RDP5 (Martin et al., 2015) – gives a putative recombination-free alignment that we call non-recombinant alignment 3 (**NRA3**) (see *Methods*).

All three approaches to removing recombinant genomic segments point to a single ancestral lineage for SARS-CoV-2 and RaTG13. Two other bat viruses (CoVZXC21 and CoVZC45) from Zhejiang province fall on this lineage as recombinants of the RaTG13/SARS-CoV-2 lineage and the clade of Hong Kong bat viruses sampled between 2005 and 2007 (Figure 1, bottom). Specifically, progenitors of the RaTG13/SARS-CoV-2 lineage appear to have recombined with the Hong Kong clade (with inferred breakpoints at 11.9kb and 20.8kb) to form the CoVZXC21/CoVZC45-lineage. Ancestors to the RaTG13/SARS-CoV-2 lineage also include a pangolin sequence sampled in Guangdong province in March 2019 and a clade of pangolin sequences from Guangxi province sampled in 2017. Major phylogenetic and phylogeographic relationships are shown in Figure 2, with phylogenies reconstructed for two major sub-regions of **NRR1**.

**Figure 2.**
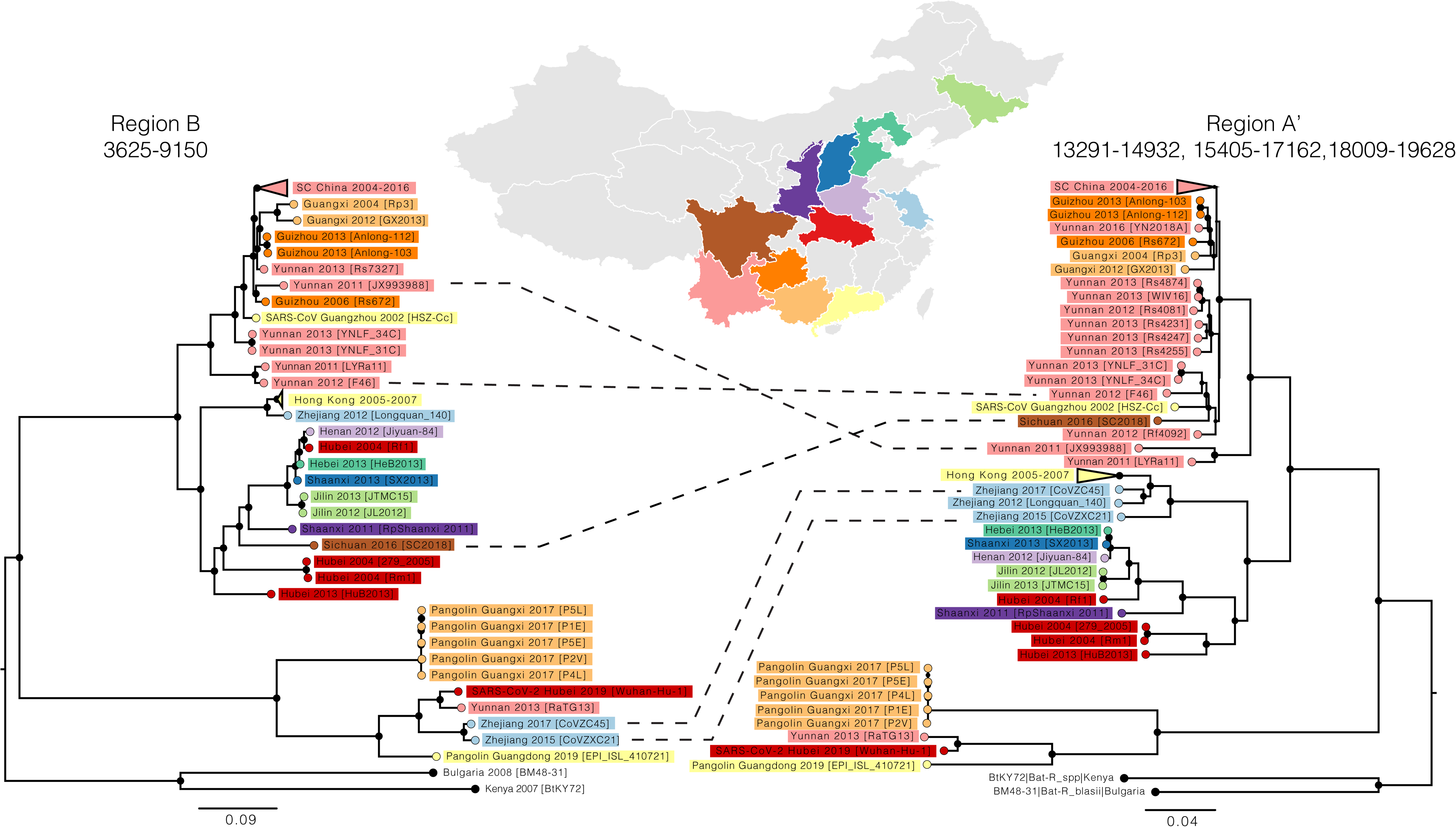
Maximum likelihood trees of the sarbecoviruses using the two longest breakpoint-free regions (BFRs), rooted on the Kenya/Bulgaria lineage. Region **A** has been shortened to **A’** (5017nt) based on potential recombination signals within the region. Region **B** is 5525nt long. Sequences are color-coded by province according to the map. Five example sequences with incongruent phylogenetic positions in the two trees are indicated by dashed lines.

As the SARS-CoV-2 S protein has been implicated in past recombination events or possibly convergent evolution (Lam et al., 2020), we specifically investigated several sub-regions of the S protein – the N-terminal domain of S1, the C-terminal domain of S1, the variable loop region of the C-terminal domain, and S2. The variable loop region in SARS-CoV-2 shows closer identity to the 2019 pangolin coronavirus sequence than to the RaTG13 bat virus, supported by phylogenetic inference (Figure 3). On first examination this would suggest that that SARS-CoV-2 is a recombinant of an ancestor of Pangolin-2019 and RaTG13. However, on closer inspection, the relative divergences in the phylogenetic tree (Figure 3, bottom) show that SARS-CoV-2 is unlikely to have acquired the variable loop from an ancestor of Pangolin-2019 as these two sequences are approximately 10% divergent throughout the entire S protein (excluding the NTD). It is RaTG13 that is divergent in the variable loop region and is the likely product of recombination, acquiring a divergent variable loop from an as yet unsampled bat sarbecovirus. This is notable because the variable loop region contains the six key contact residues in the receptor binding domain (RBD) that give SARS-CoV-2 its ACE2 binding specificity (Anderson et al., 2020; Wan et al., 2020). These residues are also in the 2019 pangolin coronavirus sequence. The most parsimonious explanation for these shared residues is a single bat virus lineage (including the common ancestor of SARS-CoV-2, RaTG13, and Pangolin Guangdong 2019) has the ACE2 specific residues, rather than the SARS-CoV-2 being recombinant. This provides compelling support for the SARS-CoV-2 lineage being the consequence of a direct or nearly-direct zoonotic jump from bats because the key ACE2 binding residues were present in viruses circulating in bats.

**Figure 3.**
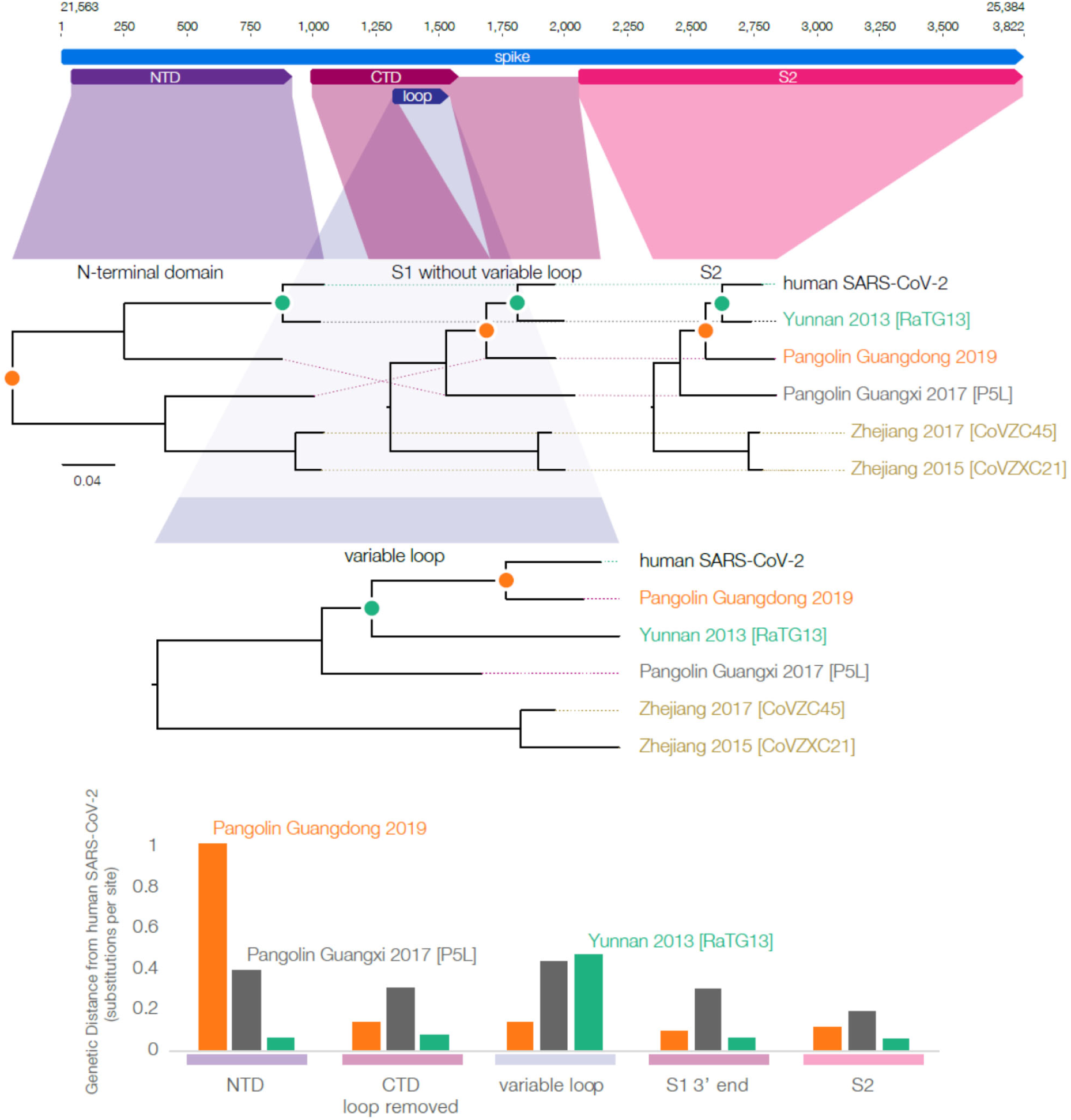
Phylogenetic relationships among SARS-CoV-2 and closely related sequences for sub-regions of the S protein. SARS-CoV-2 and RaTG13 are the most closely related (their most recent common ancestor node denoted by a green circle), except in the 220nt variable loop region of the C-terminal domain (bar graphs at bottom). In the variable loop region, RaTG13 diverges considerably with the tMRCA now outside that of SARS-CoV-2 and Pangolin Guangdong 2019 ancestor, suggesting that RaTG13 has acquired this region from a more divergent and undetected bat lineage. The genetic distances between SARS-CoV-2 and RaTG13 (bottom panel) demonstrate their relationship is consistent across all regions except for the variable loop. The genetic distances between SARS-CoV-2 and Pangolin Guangdong 2019 are consistent across all regions except the NTD.

### Ancestry in non-recombinant regions

Using the most conservative approach to identifying a non-recombinant genomic region (**NRR1**), SARS-CoV-2 forms a sister lineage with RaTG13, with genetically related cousin lineages of coronavirus sampled in pangolins in Guangdong and Guangxi provinces (Figure 2). Given that these pangolin viruses are ancestral to the progenitor of the RaTG13/SARS-CoV-2 lineage, it is more likely that they are also picking up viruses from bats. While pangolins could be acting as intermediate hosts for bat viruses to get into humans — they develop severe respiratory disease (Lie et al., 2019) and commonly come into contact with people as they are trafficked in large numbers for consumption and use in Chinese medicine — pangolin infection is not a requirement for bat viruses to cross into humans.

Phylogenies of sub-regions of **NRR1** depict an appreciable degree of spatial structuring of the sarbecoviruses population across different regions (Figure 2). One cluster consists of viruses from provinces in southcentral (SC) China (Guangxi, Yunnan, Guizhou) as well as one sequence from Sichuan province. The major cluster in its sister lineage is almost exclusively occupied by viruses from provinces in northeast (NE) and central China (Hubei, Shaanxi, Shanxi, Henan, Hebei, Zheijang and Jilin).

### tMRCA for non-recombinant regions of SARS-CoV-2 lineage

To avoid artefacts due to recombination, we focused on the non-recombining regions **NRR1**, **NRR2**, and the recombination-masked alignment **NRA3** for inferring time-measured evolutionary histories. Visual exploration using TempEst (Rambaut et al., 2016) indicates there is no evidence for temporal signal in these data sets (Figure S1). This is not surprising for diverse viral populations with relatively deep evolutionary histories. In such cases, even moderate rate variation among long deep phylogenetic branches will significantly impact expected root-to-tip divergences over a sampling time range that represents only a small fraction of the evolutionary history (Trova et al., 2015). However, formal testing using marginal likelihood estimation (Duchene et al., 2019) does not reject the absence of temporal signal in all three data sets (Table S1), albeit without strong support in favor of temporal signal (log Bayes factor support of 3, 10, and 3 for **NRR1**, **NRR2**, and **NRA3** respectively).

In the absence of a strong temporal signal, we sought to identify a suitable prior rate distribution to calibrate the time-measured trees by examining several coronaviruses sampled through time, including HCoV-OC43, MERS-CoV, and SARS-CoV virus genomes. These data sets were subjected to the same recombination masking approach as **NRA3** and were characterized by a strong temporal signal (Figure 4), but also by markedly different evolutionary rates. Specifically, using a formal Bayesian approach (Suchard et al., 2018) (see *Methods*), we estimate a fast evolutionary rate (0.00169 subst/site/yr, 95% highest posterior density (HPD) interval [0.00131,0.00205]) for SARS viruses sampled over a limited time scale (1 year), a slower rate (0.00078 [0.00063,0.00092] subst/site/yr) for MERS-CoV on a time scale of about 4 years, and the slowest rate (0.00024 [0.00019,0.00029] subst/site/yr) for HCoV-OC43 over almost five decades. These differences reflect the fact that rate estimates can vary considerably with the time scale of measurement, a frequently observed phenomenon in viruses known as time-dependent evolutionary rates (Aiewsakun and Katzourakis, 2016; Duchene et al., 2014; Membrebe et al., 2019). Over relatively shallow time-scales, such differences can primarily be explained by varying selective pressure with mildly deleterious variants being eliminated more strongly by purifying selection over longer time scales (Holmes, 2009; Holmes et al., 2016; Membrebe et al., 2019). In line with this, we estimate a concomitantly decreasing nonsynonymous-to-synonymous substitution rate ratio (dN/dS) over longer evolutionary time-scales: 1.41 [1.20,1.68], 0.35 [0.30,0.41], and 0.133 [0.129,0.136] for SARS, MERS-CoV, and HCoV-OC43 respectively. In the light of these time-dependent evolutionary rate dynamics, a rate in the slow range is appropriate for calibrating the sarbecovirus evolutionary history, but we compare both MERS-CoV and HCoV-OC43 centred prior distributions (Figure S2) with relatively large variances in our subsequent analyses in order to examine the sensitivity of the date estimates to this prior specification.

**Figure 4.**
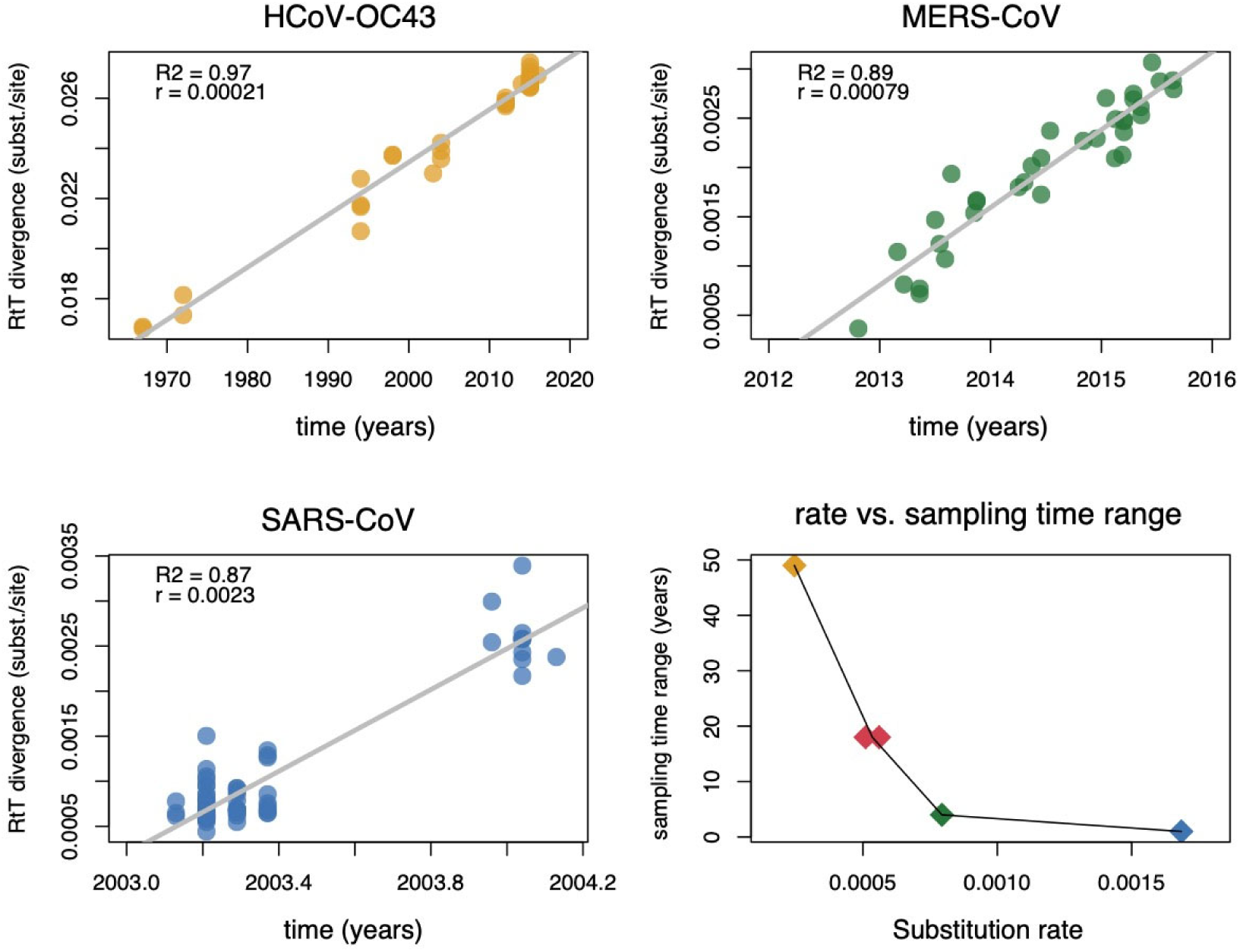
Temporal signal and mean evolutionary rate estimates for HCoV-OC43, MERS, and SARS coronaviruses. (**A-C**) Root-to-tip divergence as a function of sampling time for the three coronavirus evolutionary histories unfolding over different time scales. (**D**) Mean evolutionary rate estimates plotted against sampling time range for the same three data sets (represented by the same color as the data points in their respective root-to-tip divergence plots), as well as for the comparable recombination-masked sarbecovirus alignment (**NRA3**) using the two different priors for the rate in the Bayesian inference (red points).

We infer time-measured evolutionary histories using a Bayesian phylogenetic approach while incorporating rate priors based on the mean MERS-CoV and HCoV-OC43 rates and with standard deviations that allow for more uncertainty than the empirical estimates for both viruses (see *Methods*). Using both prior distributions, this results in six highly similar posterior rate estimates for **NRR1**, **NRR2**, and **NRA3**, centered around 0.00055 subst/site/yr. The fact that these estimates lie in between the MERS-CoV and HCoV-OC43 rate is consistent with the intermediate sampling time range of about 18 years (Figure 5). The consistency of the posterior rates for the different prior means also implies that the data do contribute to the evolutionary rate estimate despite the fact that a temporal signal was visually not apparent (Figure S1). Below, we report divergence time estimates based on the HCoV-OC43-centred rate priors, but also summarize similar corresponding estimates for the MERS-CoV-centred rate priors in Supplementary Figure S3.

**Figure 5.**
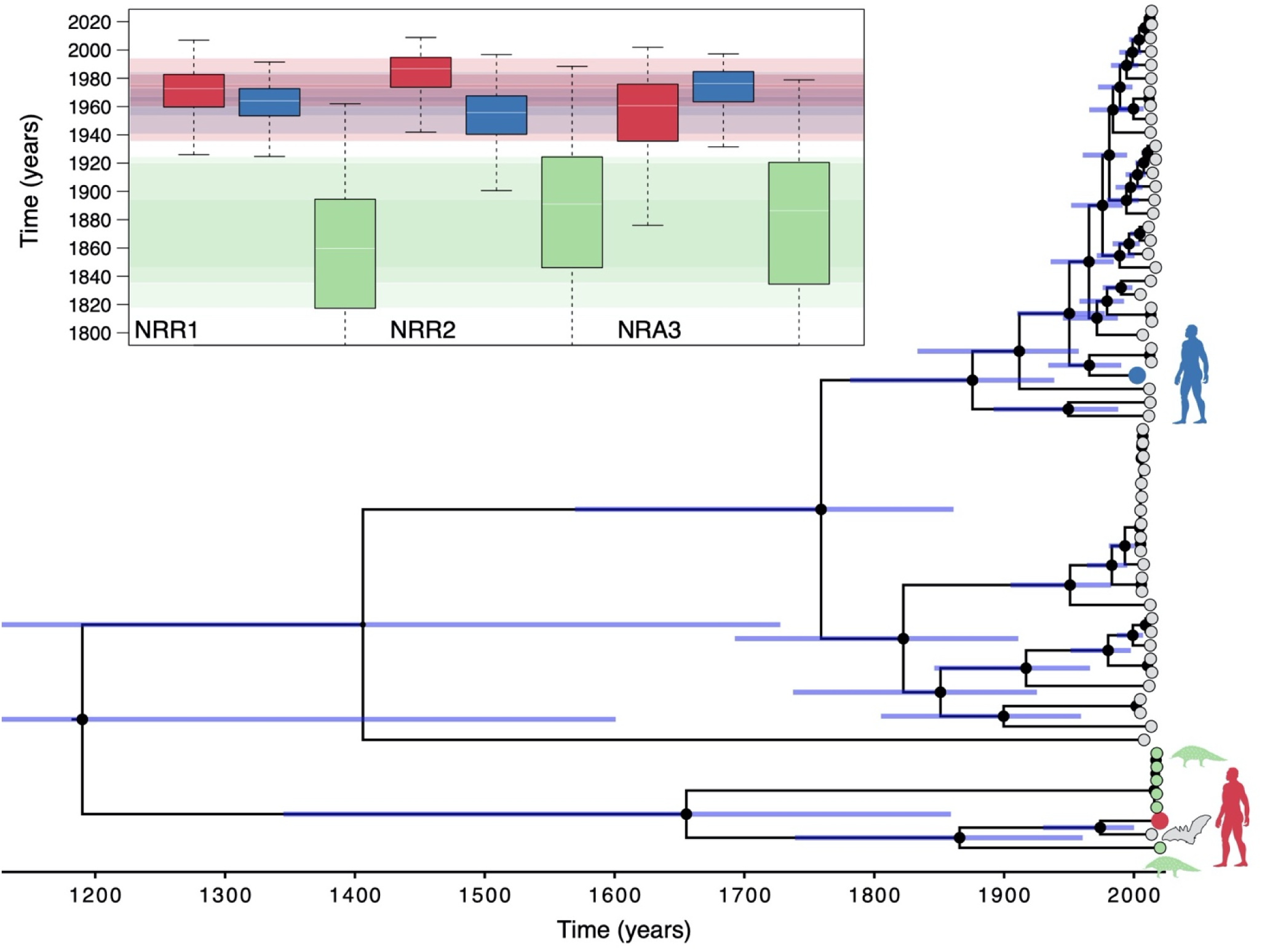
Time-measured phylogenetic estimates and divergence times for sarbecovirus lineages using an HCoV-OC43 centred rate prior. The time-calibrated phylogeny represents a maximum clade credibility tree inferred for **NRR1**. Grey tips correspond to bat viruses, green to pangolin, blue to the SARS-CoV virus, and red to SARS-CoV-2. The size of the black internal node circles are proportional to the posterior node support. 95% credible intervals bars are shown for all internal node ages. The inset represents divergence time estimates based on **NRR1**, **NRR2** and **NRA3**. The boxplots represent the divergence time estimates for SARS-CoV-2 (red boxplot) and the 2002-2003 SARS-CoV virus (blue boxplot) from their most closely related bat virus. Green boxplots show the TMRCA estimate for the RaTG13/SARS-CoV-2 lineage and its most closely related pangolin lineage (Guangdong 2020). Transparent boxes of the interquartile range width and with the same colours are superimposed to highlight the overlap between the estimates. In Supplementary Figure S3, we compare these divergence time estimates to those obtained using the MERS-CoV-centred rate priors for **NRR1**, **NRR2** and **NRA3**.

The divergence time estimates for SARS-CoV-2 and SARS-CoV from their respective most closely related bat lineages are reasonably consistent among the three approaches we use to eliminate the effects of recombination in the alignment. Using the most conservative approach (**NRR1**), the divergence time estimate for SARS-CoV-2 and RaTG13 is 1969 (95% HPD: 1930-2000) while the divergence time estimate between SARS-CoV and its most closely related bat sequence is 1962 (95% HPD: 1932-1988); see Figure 5. These are in general agreement with estimates using **NRR2** and **NRA3**, which result in divergence times of 1982 [1948,2009] and 1948 [1879-1999], respectively, for SARS-CoV-2, and estimates of 1952 [1906, 1989] and 1970 [1932-1996], respectively, for the divergence time of SARS-CoV from its closest known bat ancestor. The SARS-CoV divergence times are somewhat earlier than previously estimated dates (Hon et al., 2008) as previous estimates were obtained using a collection of SARS-CoV genomes from human and civet hosts (as well as a few closely related bat genomes) which implies that the evolutionary rates were predominantly informed by the short-term SARS outbreak scale and likely biased upwards for the time scale under investigation. Indeed, the rates reported by these studies are in line with the short-term SARS rates we estimate (Figure 4). The estimated divergence times for the pangolin virus most closely related to SARS-CoV-2/RaTG13 range from 1851 [1730,1958] to 1877 [1746,1986], indicating that these pangolin lineages were acquired from bat viruses divergent to those that gave rise to SARS-CoV-2. Current sampling of pangolins does not implicate them as an intermediate host.

## Discussion

Identifying the origins of an emerging pathogen can be critical during the early stages of an outbreak as it may allow for containment measures to be precisely targeted at a stage when the number of daily new infections is still low. Early detection via genomics and sequencing was not possible during SE Asia’s initial outbreaks of avian influenza H5N1 (1997, 2003-2004) or the first SARS outbreak (2002-2003). By 2009 however rapid genomic analysis was a routine component of outbreak response. The 2009 influenza pandemic and subsequent outbreaks of MERS-CoV (2012), H7N9 avian influenza (2013), Ebola virus (2014), and Zika virus (2015) were met with rapid sequencing and genomic characterization. For the current pandemic, the ‘novel pathogen identification’ component of outbreak response delivered on its promise, with viral identification and rapid genomic analysis providing confirmation and a genome sequence, within weeks, that the December 2019 outbreak in Wuhan was caused by a coronavirus (World Health Organization, 2020a). Unfortunately, a response that would achieve containment was not possible. Given what was known about the origins of SARS as well as identification of SARS-like viruses circulating in bats that had binding sites adapted to human receptors (Ge et al., 2013; Menachery et al., 2015, 2016), appropriate measures should have been in place for outbreaks of novel coronaviruses. The key to successful surveillance is that we know which human-adapted viral phenotypes to look for (Holmes et al., 2018).

The key difficulty in inferring reliable evolutionary histories for coronaviruses is that their high recombination rate (Graham and Baric, 2010; Su et al., 2016) violates the assumption of standard phylogenetic approaches because different parts of the genome will have different origins. In order to begin characterizing any ancestral relationships for SARS-CoV-2, non-recombinant regions of the genome must be identified, so that reliable phylogenetic reconstruction and dating can be performed. Evolutionary rate estimation can be profoundly affected by the presence of recombination (Schierup and Hein, 1999). As there is no single accepted method of inferring breakpoints and identifying clean sub-regions with high certainty, we implemented several approaches to identifying three classic statistical signals of recombination: mosaicism, phylogenetic incongruence and excessive homoplasy (Posada et al., 2002). Our most conservative approach attempted to ensure that putative non-recombining regions had no mosaic or phylogenetic incongruence signals. A second approach was conservative with respect to breakpoint identification, while a third approach attempted to minimize the number of regions removed while still minimizing signals of mosaicism and homoplasy. The origins we present in Figure 5 (**NRR1**) are conservative in the sense that **NRR1** is more likely to be non-recombinant than **NRR2** or **NRA3**. As the estimated rates and divergence dates were highly similar in the three data sets analyzed, we conclude that our estimates are robust to the method of identifying the genome’s non-recombinant regions.

Due to the absence of temporal signal in the sarbecovirus data sets, we used informative prior distributions on the evolutionary rate to estimate divergence dates. We show that such prior calibration information can be obtained from other coronaviruses (SARS-CoV, MERS-CoV, and HCoV-OC43), but their rates vary with the time scale of sample collection. In the presence of time-dependent rate variation, a widely observed phenomenon for viruses (Duchene et al., 2014; Aiewsakun and Katzourakis, 2016; Membrebe et al., 2019), the slower rates appear more appropriate for sarbecoviruses that currently encompass a sampling time range of about 18 years. We therefore centered the prior rate distributions on the mean HCoV-OC43 and MERS-CoV rate, but permitted the distributions a greater variance than the empirical rate estimates for these lineages. This approach resulted in similar posterior rates despite the different prior means, implying that the sarbecovirus data do inform the rate estimate even though root-to-top temporal signal was not apparent.

The relatively fast evolutionary rate means that it is most appropriate to estimate shallow nodes in the sarbecovirus evolutionary history including the divergence times of SARS-CoV and SARS-CoV-2 from their most closely related bat viruses. Accurately estimating deeper nodes would require adequately accommodating time-dependent rate variation. While such models have recently been made available, we lack the information to calibrate the rate decline through time (e.g. through internal node calibrations (Membrebe et al., 2019)). As a proxy, it would be possible to model the long-term purifying selection dynamics as a major source of time-dependent rates rates (Duchene et al., 2014; Aiewsakun and Katzourakis, 2016; Membrebe et al., 2019), but this is beyond the scope of the current study. The assumption of long-term purifying selection would imply that coronaviruses are at endemic equilibrium with their natural host species, horseshoe bats, to which they are presumably well-adapted. Although there is currently little evidence supporting or contradicting strong positive selection in the sarbecovirus lineage, if multiple host species were to be identified that are able to support endemic viral transmission, then the positive selection that would be associated with new species adaptations would need to be considered when inferring evolutionary rate variation through time.

Of importance for future emergence events is the appreciation that SARS-CoV-2 has emerged from the same horseshoe bat subgenus that harbours SARS-like coronaviruses. Another similarity between SARS-CoV and SARS-CoV-2 is their divergence time (40-70 years ago) from currently known extant bat-virus lineages (Figure 5). This long divergence period indicates there are unsampled viruses circulating in horseshoe bats that have zoonotic potential (Zhou et al., 2020a). While there is clear involvement of other mammalian species – specifically pangolins for SARS-CoV-2 – as a plausible conduit for transmission to humans, there is no evidence pangolins are facilitating adaptation to humans. A hypothesis of snakes as intermediate hosts of SARS-CoV-2 was posited during the early epidemic phase (Ji et al., 2020), but we found no evidence of this (Anderson, 2020; Robertson, 2020); see Supplementary Section 3.

With horseshoe bats currently the most plausible source for SARS-CoV-2, it is important to consider that sarbecoviruses circulate in a variety of horseshoe bat species with often widely overlapping species ranges (Wong et al., 2019). Yet, the viral population is largely spatially structured according to provinces in the south and southeast on one lineage, and provinces in the center, east, and northeast on another (Figure 2). This boundary appears to be rarely crossed. Two exceptions can be seen in the relatively close relationship of viruses from Hong Kong bats to viruses from Zheijang (with two of the latter, CoVZC45 and CoVZXC21, identified as recombinants) and a recombinant virus from Sichuan for which part of the genome (region B of SC2018 in Figure 2) clusters with viruses from provinces in the center, east, and northeast of China. SARS-CoV-2 and RaTG13 are also exceptions as they were sampled from Hubei and Yunnan, respectively. The fact that they are geographically relatively distant is in agreement with their somewhat distant tMRCA because the spatial structure suggests that migration between their locations may be uncommon. From this perspective, it may be useful to perform surveillance for more closely related virus to SARS-CoV-2 along the gradient from Yunnan to Hubei.

It is clear from our analysis that viruses closely related to SARS-CoV-2 have been circulating in horseshoe bats for many decades. The substantial unsampled diversity on the SARS-CoV-2/RaTG13 lineage suggests that there is a major clade of bat sarbecoviruses with generalist properties – with respect to their ability to infect a range of mammalian cells – that facilitated its jump to humans and may do so again. Although the human ACE2-compatible receptor binding domain was very likely to have been present in a bat sarbecovirus lineage that ultimately lead to SARS-CoV-2, this RBD sequence has thus far only been found in a few pangolin viruses. Furthermore, the other key feature thought to be instrumental to SARS-CoV-2’s ability to infect humans – a polybasic cleavage site insertion in the Spike protein – has been seen in another close bat relative of the SARS-CoV-2 virus (Zhou et al., 2020a); however, the sequence is different and it is likely an independent event.

The existing diversity and dynamic process of recombination amongst lineages in the bat reservoir demonstrate how difficult it will be to identify viruses with potential to cause significant human outbreaks before they emerge. This underscores the need for a global network of real-time human disease surveillance systems like that which identified the unusual cluster of pneumonia in Wuhan in December 2019 and the rapid deployment of functional studies and genomic tools for pathogen identification and characterization.

## Methods

### Data set compilation

Sarbecovirus data. Complete genome sequence data were downloaded from GenBank and ViPIR; accession numbers available in Supplementary Section 4. Sequences were aligned by MAFTT (Katoh et al., 2009), with a final alignment length of 30927, and used in the analyses below.

HCoV-OC43. We compiled a data set including 27 human coronavirus OC43 virus genomes and 10 related animal virus genomes (6 bovine, 3 white-tailed deer and one canine virus). The canine viral genome was excluded from the Bayesian phylogenetic analyses because temporal signal analyses (see below) indicated that it was an outlier.

MERS-CoV. We extracted a similar number (*n* = 35) of genomes from a MERS-CoV data set analysed by Dudas et al (2018) using the phylogenetic diversity analyser tool (Chernomor et al., 2015).

SARS-CoV. We compiled a set of 69 SARS-CoV genomes including 58 genomes sampled from human and 11 sampled from civets and raccoon dogs. This data set comprises an updated version of the one used in Hon et al (2008).

### Recombination analysis

As coronaviruses are known to be highly recombinant, we used three different approaches to identify non-recombinant regions to be used in our Bayesian time-calibrated phylogenetic inference.

First, we took an approach that relies on identifying mosaic regions (via 3SEQ) that are also supported by phylogenetic incongruence (PI) signals (Boni et al., 2010). As 3SEQ is the most powerful the mosaic methods (Boni et al., 2007), we used 3SEQ to identify the best supported breakpoint history for each potential child (recombinant) sequence in the data set. A single 3SEQ run on the genome alignment resulted in 67 out of 68 sequences supporting some recombination in the past, with multiple candidate breakpoint ranges listed for each putative recombinant. Then, we (a) collected all breakpoints into a single set, (b) complemented this set to generate a set of non-breakpoints, (c) grouped the non-breakpoints into contiguous breakpoint-free regions (BFRs), and (d) sorted these regions by length. A phylogenetic tree – using RAxML v8.2.8 (Stamatakis, 2014, 2006), GTR+Γ model, 100 bootstrap replicates – was inferred for each BFR longer than 500nt.

We considered both (1) the possibility that BFRs could be combined into larger non-recombinant regions, and (2) the possibility of further recombination within each BFR.

We named the length-sorted BFRs as: region **A** (nt positions 13291-19628, length=6338nt), region **B** (nt positions 3625-9150, length=5526nt), region **C** (nt positions 9261-11795, length=2535nt), region **D** (nt positions 27702-28843, length=1142nt), and six more through region **J**. Phylogenetic trees for all ten BFRs are shown in Supplementary Figures S5-S14. Regions **A**, **B**, and **C** had similar phylogenetic relationships among the southcentral China bat viruses (Yunnan, Guangxi, Guizhou provinces), the Hong Kong viruses, northeastern Chinese viruses (Jilin, Shaanxi, Shanxi, Hebei, Henan provinces), pangolin viruses, and the SARS-CoV-2 lineage. As these subclades had different phylogenetic relationships in region **D** (Supplementary Section 5), region **D** and shorter BFRs were not further included into a combined putative non-recombinant regions.

Regions **A**, **B**, and **C** were further examined for mosaic signals by 3SEQ; all showed signs of mosaicism. In region **A**, we removed sub-region **A1** (nt positions 3872-4716 within region **A**) and sub-region **A4** (nt 1642-2113) as both of these showed PI-signals with other sub-regions of region **A**. After removal of **A1** and **A4**, we named the new region **A′**. In addition, sequences NC_014470 (Bulgaria 2008), CoVZXC21, CoVZC45, and DQ412042 (Hubei-Yichang) needed to be removed to maintain a clean non-recombinant signal in **A′**. Region **B** showed no PI-signals within the region, except one including sequence SC2018 (Sichuan), thus this sequence was also removed from the set. Region **C** showed no PI-signals within it. Combining regions **A′**, **B**, and **C**, and removing the five mentioned sequences gives us putative non-recombining region 1, or **NRR1**, as an alignment of 63 sequences.

Second, we wanted to construct non-recombinant regions where our approach to breakpoint identification was as conservative as possible. In this approach, we considered a breakpoint supported only if it had three types of statistical support from (1) mosaic signals identified by 3SEQ, (2) PI-signals identified by building trees around 3SEQ’s breakpoints, (3) the GARD algorithm (Kosakovsky Pond et al., 2006) which identifies breakpoints by identifying PI-signals across proposed breakpoints. Since 3SEQ identified 10 BFRs longer than 500nt, we used GARD’s inference on 10, 11, and 12 breakpoints. A reduced sequence set of 25 sequences chosen to capture the breadth of diversity in the sarbecoviruses (obvious recombinants not involving the SARS-CoV-2 lineage were also excluded) was used as GARD is computationally intensive. GARD identified eight breakpoints that were also within 50nt of breakpoints identified by 3SEQ. PI-signals were identified (with bootstrap support >80%) for seven of these eight breakpoints: positions 1684, 3046, 9237, 11885, 21753, 22773, and 24628. Using these breakpoints, the longest putative non-recombining segment (nt 11885-21753) is 9.9kb long, and we call this region **NRR2**. Because this approach to breakpoint identification is conservative, the approach to identifying non-recombinant regions is not conservative.

Our third approach involved identifying breakpoints and masking minor recombinant regions (with gaps, which are treated as unobserved characters in probabilistic phylogenetic approaches). Specifically, we used a combination of six methods implemented in RDP5 (RDP, GENECONV, MaxChi, Bootscan SisScan and 3SEQ) and considered recombination signals detected by more than two methods for breakpoint identification. Except for specifying that sequences are linear, all settings were kept to their defaults. Based on the identified breakpoints in each genome, only the major non-recombinant region is kept in each genome while other regions are masked. To evaluate the performance procedure, we confirmed that the recombination masking resulted in (*i*) a significantly different outcome of the PHI-test (Bruen et al., 2006), (*ii*) a removal of well-supported (bootstrap value >95%) incompatible splits in Neighbor-Nets (Bryant and Moulton, 2004), and (*iii*) a near-complete reduction of mosaic signal as identified by 3SEQ. If the latter still identified non-negligible recombination signal, we removed additional genomes that were identified as the major contributors to the remaining signal. This produced non-recombining alignment **NRA3**, which included 63 of the 68 genomes.

### Bayesian divergence time estimation

We focused on these three non-recombining regions/alignments for divergence time estimation. This avoids inappropriately modelling evolutionary processes with recombination on strictly bifurcating trees, which can result in different artefacts such as homoplasies that inflate branch lengths and result in apparently longer evolutionary divergence times. In order to examine temporal signal in the sequenced data, we plotted root-to-tip divergence against sampling time using TempEst (Rambaut et al., 2016) based on a maximum likelihood tree. The latter was reconstructed using IQTREE (Nguyen et al., 2014) under a General-Time Reversible (GTR) model with a discrete gamma distribution to model among-site rate variation.

Time-measured phylogenetic reconstruction was performed using a Bayesian approach implemented in BEAST (Suchard et al., 2018). When the genomic data included both coding and noncoding regions we used a single GTR+Γ substitution model; for concatenated coding genes we partitioned the alignment by codon positions and specified an independent GTR+Γ model for each partition with a separate gamma model to accommodate among-site rate variation. We used an uncorrelated relaxed clock model with a lognormal distribution for all data sets, except for the low-diversity SARS data for which we specified a strict molecular clock model. For the HCoV-OC43, MERS-CoV, and SARS data sets we specified flexible skygrid coalescent tree priors. In the absence of any reasonable prior knowledge on the tMRCA of the sarbecovirus data sets (which is required for the grid specification in a skygrid model), we specified a simpler constant size population prior. As informative rate priors for the analysis of the sarbecovirus data sets, we used two different normal prior distributions: one with a mean of 0.00078 and a standard deviation of 0.0003 and one with a mean of 0.00024 and standard deviation of 0.0001. These means are based on the mean rates estimated for MERS-CoV and HCoV-OC43 respectively, while the standard deviations are set ten times larger than the empirical standard deviations to allow more prior uncertainty and avoid strong bias (Supplementary Fig. 2). In our analyses of the sarbecovirus data sets, we incorporated the uncertainty of the sampling dates when exact dates were not available. To estimate nonsynonymous over synonymous rate ratios for the concatenated coding genes, we used the empirical Bayes ‘Renaissance counting’ procedure (Lemey et al., 2012). Temporal signal was tested using a recently developed marginal likelihood estimation procedure (Duchene et al., 2019) (Table S1).

Posterior distributions were approximated through Markov chain Monte Carlo sampling, which were run sufficiently long to ensure effective sampling sizes > 100. BEAST inferences made use of the BEAGLEv3 library (Yres et al., 2019) for efficient likelihood computations. We used TreeAnnotator to summarize the posterior tree distributions and annotated the estimated to a maximum clade credibility tree, which was visualized using FigTree.

## Author Contributions

All authors contributed to analyses and interpretations. DLR and XJ performed recombination and phylogenetic analysis, and annotated the virus names with geographical and sampling dates. AR performed the S recombination analysis. PB and TC performed the codon-usage analysis. MFB performed the recombination analysis for non-recombining regions 1 and 2, breakpoint analysis, and phylogenetic inference on recombinant segments. PL performed recombination analysis for non-recombining alignment 3, calibration of rate of evolution, and phylogenetic reconstruction and dating. TT-YL collected SARS-CoV data. PL, MFB, and DLR wrote the first draft of the manuscript, and all authors contributed to manuscript editing.

## Acknowledgements

We would like to thank all the authors who have kindly deposited and shared genome data on GISAID. Thanks to Trevor Bedford for providing MFB an alignment on which an initial recombination analysis was done. The research leading to these results has received funding from the European Research Council under the European Union’s Horizon 2020 research and innovation programme (grant agreement no. 725422-ReservoirDOCS). DLR is funded by the MRC (MC UU 1201412). The Artic Network receives funding from the Wellcome Trust through project 206298/Z/17/Z. PL acknowledges support by the Research Foundation -- Flanders (‘Fonds voor Wetenschappelijk Onderzoek -- Vlaanderen’, G066215N, G0D5117N and G0B9317N). TL is funded by The National Natural Science Foundation of China (NSFC) Excellent Young Scientists Fund (Hong Kong and Macau) (31922087).

