## Supplementary Materials and Figures for "Evolutionary origins of the SARS-CoV-2 sarbecovirus lineage responsible for the COVID-19 pandemic"

### 1. Assessing temporal signals using TempEst and BETS

Root-to-tip divergence plots as a function of sampling time indicate no clear pattern of divergence accumulation over the sampling time range for all three data sets (Fig. S1).

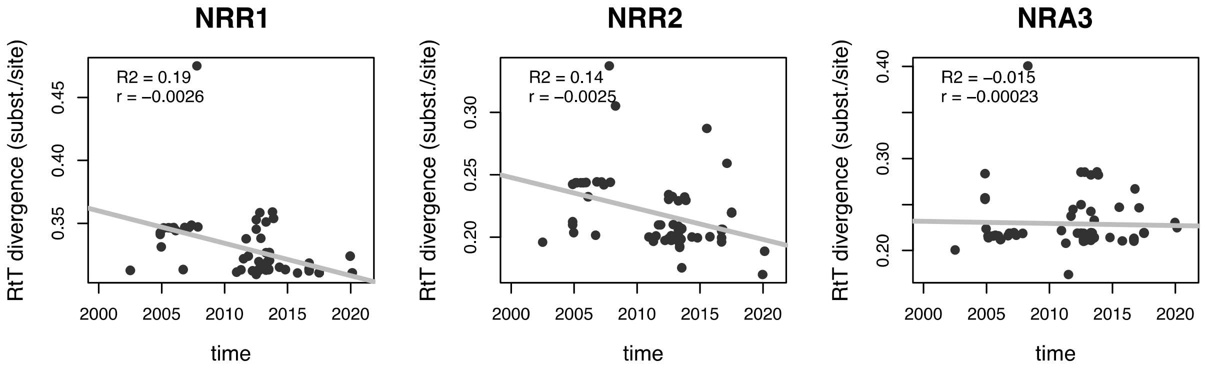


**Figure S1.** Root-to-tip divergence as a function of sampling time for non-recombinant regions **NRR1** & **NRR2** and recombination-masked alignment set (**NRA3**). The plots are based on maximum likelihood tree reconstructions with a root position that maximises the residual mean squared for the regression of root-to-tip divergence and sampling time.

To formally test for temporal signal, we used a recently proposed Bayesian model testing approach that compares the marginal likelihoods estimated for a model that constrains tips in time-measured trees to be proportional to their sampling times and has an estimable evolutionary rate to a model that enforces an ultrametric tree (all sequences sampled at the same time, reflecting that sampling times do not predict tip positions) without a free evolutionary rate parameter (Duchene et al. 2019) .  Table S1 reports the marginal likelihood estimates for these models fitted to both data sets. This results in a log Bayes factor (log BF) support of 3, 10 and 3 in favour of temporal signal for **NRR1**, **NRR2,** and **NRA3**.

**Table S1**: Log marginal likelihood estimates for a dated tip model versus an ultrametric model for **NRR1**, **NRR2** and **NRA2**.

| Data set | Log MLE dated tips | Log MLE ultrametric |
|---|---|---|
| **NRR1** | -91357 | -91360 |
| **NRR2** | -71620 | -71630 |
| **NRA2** | -118921 | -118924 |

### 2. Estimating evolutionary rates and divergence dates for NRR1, NRR2, and NRA3

Although formal testing does not reject the presence of temporal signal, the support for this signal remains limited. We therefore aimed to take advantage of prior information on the evolutionary rate in the Bayesian analysis of the three data sets, but in a way that avoids having to make strong assumptions on whether sarbecovirus evolutionary rates should closely match the MERS-CoV or HCoV-OC43 rates. We adopt two evolutionary rate priors that are centred on the mean rates for both MERS-CoV and HCoV-OC43, but with standard deviations that are ten times larger than the posterior rate distributions for MERS-CoV and HCoV-OC43 (Fig. S2). Using these priors, we infer highly similar evolutionary rate posteriors for **NRR1**, **NRR2,** and **NRA3**. In addition, the posterior rate distributions show a considerable reduction in variance compared to their priors (Fig. S2), indicating that sampling dates provide information about the rate despite the difficulties in ascertaining the temporal signal based on visual exploration.

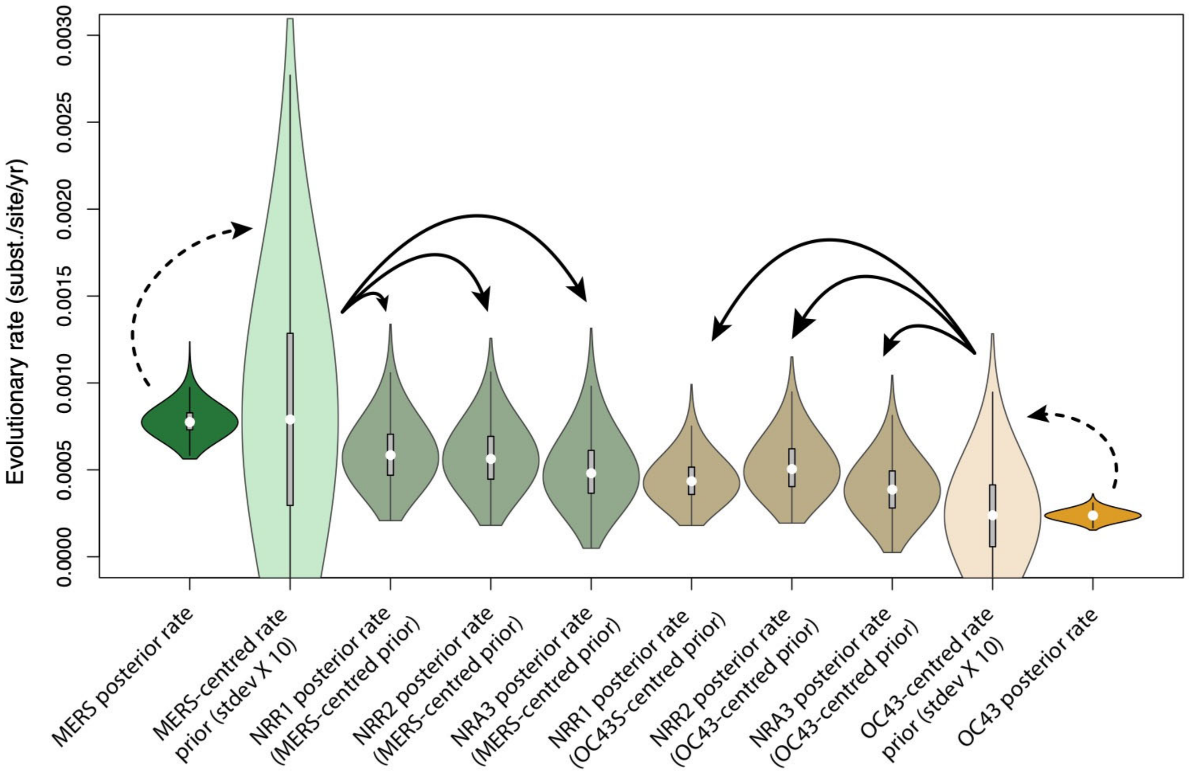


**Figure S2.** Posterior rate distributions for MERS-CoV (far left) and HCoV-OC43 (far right) using BEAST on *n*=27 sequences spread over 4 years (MERS-CoV) and *n*=27 sequences spread over 49 years (HCoV-OC43). As illustrated by the dashed arrows, these two posteriors motivate (dashed arrows) our specification of prior distributions with standard deviations inflated 10-fold (light color). These rate priors are subsequently used in the Bayesian inference of posterior rates for **NRR1, NRR2,** and **NRA3** as indicated by the solid arrows.

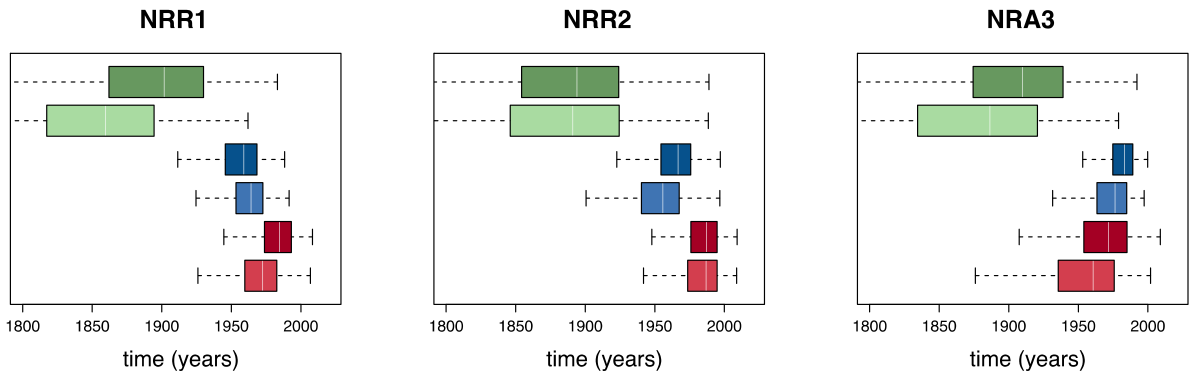


**Figure S3.** Divergence time estimates based on the three regions/alignments where the effects of recombination have been removed. The red and blue boxplots represent the divergence time estimates for SARS-CoV-2 (red) and the 2002-2003 SARS-CoV (blue) from their most closely related bat virus, with the light- and dark-colored versions based on the HCoV-OC43 and MERS-CoV centered priors, respectively. Green boxplots show the TMRCA estimate for the RaTG13/SARS-CoV-2 lineage and its most closely related pangolin lineage (Guangdong 2019), with the light and dark coloured version based on the HCoV-OC43 and MERS-CoV centred priors, respectively. TMRCA estimates for SARS-CoV-2 and SARS-CoV from their respective most closely related bat lineages are reasonably consistent for the different data sets and different rate priors in our analyses. The mean estimates with 95% HPDs are listed in Table S2.

**Table S2.** Posterior estimates for the divergence time of RaTG13 bat virus and SARS-CoV-2

| | HCoV-OC43 rate prior | MERS-CoV rate prior |
|---|---|---|
| NRR1 | 1969 (95% HPD: 1930-2000) | 1981 (95% HPD: 1951-2007) |
| NRR2 | 1982 (95% HPD:1948-2009) | 1984 (95% HPD: 1953-2007) |
| NRA3 | 1948 (95% HPD:1879-1999) | 1966 (95% HPD: 1914-2002) |

**3. SARS-CoV-2 synonymous codon usage patterns are not consistent with snake genome**

Our analysis of SARS-CoV-2 origins would not be complete without mentioning why the widespread reporting of the ostensibly unique part of the SARS-CoV-2 lineage was recombinant involving a snake virus (Ji et al. 2020). To demonstrate why this is the case a heat map of the relative synonymous codon usage (RSCU) metric for this expanded dataset revealed four distinct clusters of species consisting of (1) all coronaviruses, (2) invertebrates (low GC), (3) vertebrates with high GC content, and (4) vertebrates with low GC content (Figure S4a). The cluster comprised of coronaviruses was most similar to the cluster of invertebrates, which possess the lowest average GC content of all eukaryotes analyzed. Notably, the cluster of coronaviruses is no more similar to the cluster containing snakes than it is to the cluster containing vertebrates with higher GC content that includes the representative bat species. The first two principle components of a PCA of RSCU values for all species clearly distinguished between coronaviruses and eukaryotes, with no clustering between SARS-CoV-2 and any snake species or the five bat-derived coronaviruses with the representative bat species (Figure S4b). Additionally, a PCA of only snakes, bat, and coronavirus RSCU measures further illustrate that all sampled coronaviruses generally cluster more closely with snakes than they do with the bat, despite some of them having known bat-derived origins (Figure S4c). Lastly, squared Euclidian distance measures of RSCU between 2019-nCoV and all other species is linearly correlated with GC content, such that species with a lower GC content similar to that of 2019-nCoV exhibit a more similar profile of RSCU than do species with higher GC content (Figure S4d), suggesting that Ji et al.'s finding that snakes had the most similar RSCU to SARS-CoV-2 is due simply to the fact that snakes possess the lowest GC content of species that they analyzed, rather than snakes being a likely host reservoir of the coronavirus.

**Figure S4.** Conducting analogous analyses of codon usage bias as Ji et al. (2020) with additional (and higher quality) snake coding sequence data and several miscellaneous eukaryotes with low genomic GC content failed to find any meaningful clustering of the SARS-CoV-2 with snake genomes (A). Instead, similarity in codon usage metrics between the SARS-CoV-2 and eukaryotes analyzed was correlated with coding sequence GC content of the eukaryote, with more similar codon usage being identified in eukaryotes with low GC content similar to that of the coronavirus (B).

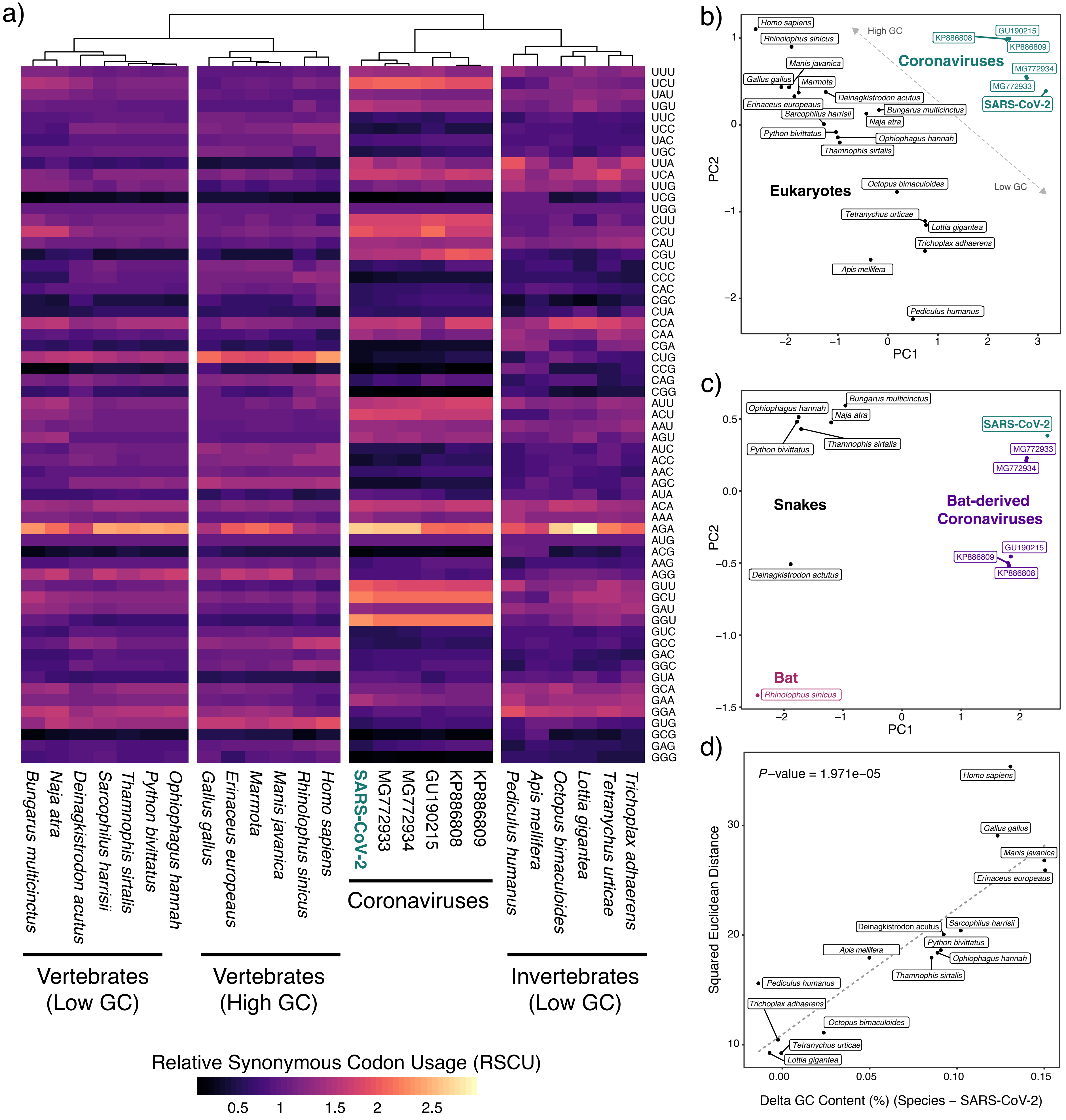

a)
UUU
UCU
UAU
UGU
UUC
UCC
UAC
UGC
UUA
UCA
UUG
UCG
UGG
CUU
CCU
CAU
CGU
CUC
CCC
CAC
CGC
CUA
CCA
CAA
CGA
CUG
CCG
CAG
CGG
AUU
ACU
AAU
AGU
AUC
ACC
AAC
AGC
AUA
ACA
AAA
AGA
AUG
ACG
AAG
AGG
GUU
GCU
GAU
GGU
GUC
GCC
GAC
GGC
GUA
GCA
GAA
GGA
GUG
GCG
GAG
GGG
Bungarus multicinctus
Naja atra
Deinagkistrodon acutus
Sarcophilus harrisii
Thamnophis sirtalis
Python bivittatus
Ophiophagus hannah
Gallus gallus
Erinaceus europeaus
Marmota
Manis javanica
Rhinolophus sinicus
Homo sapiens
SARS-CoV-2
MG772933
MG772934
GU190215
KP886808
KP886809
Pediculus humanus
Apis mellifera
Octopus bimaculoides
Lottia gigantea
Tetranychus urticae
Trichoplax adhaerens
Coronaviruses
Vertebrates (Low GC)
Vertebrates (High GC)
Invertebrates (Low GC)
Relative Synonymous Codon Usage (RSCU)
0.5
1
1.5
2
2.5
b)
High GC
Low GC
Coronaviruses
Eukaryotes
Homo sapiens
Rhinolophus sinicus
Manis javanica
Gallus gallus
Marmota
Deinagkistrodon acutus
Erinaceus europeaus
Sarcophilus harrisii
Bungarus multicinctus
Naja atra
Python bivittatus
Ophiophagus hannah
Thamnophis sirtalis
Octopus bimaculoides
Tetranychus urticae
Lottia gigantea
Trichoplax adhaerens
Apis mellifera
Pediculus humanus
KP886808
GU190215
KP886809
MG772934
MG772933
SARS-CoV-2
PC1
PC2
c)
Snakes
Bat-derived Coronaviruses
Bat
Bungarus multicinctus
Ophiophagus hannah
Naja atra
Python bivittatus
Thamnophis sirtalis
Deinagkistrodon actutus
Rhinolophus sinicus
SARS-CoV-2
MG772933
MG772934
GU190215
KP886809
KP886808
PC1
PC2
d)
P–value = 1.971e–05
Homo sapiens
Gallus gallus
Manis javanica
Erinaceus europeaus
Deinagkistrodon acutus
Sarcophilus harrisii
Apis mellifera
Python bivittatus
Ophiophagus hannah
Thamnophis sirtalis
Pediculus humanus
Trichoplax adhaerens
Octopus bimaculoides
Tetranychus urticae
Lottia gigantea
Squared Euclidean Distance
Delta GC Content (%) (Species – SARS-CoV-2)

#### 3.1. Methods for codon usage analysis

*Synonymous codon usage patterns*

Coding sequences were downloaded for 4 additional snake species (*Ophiophagus hannah* (GenBank: GCA_000516915.1) (Vonk et al. 2013), *Python bivittatus* (GenBank: GCA_000186305.2) (Castoe et al. 2013), *Thamnophis sirtalis* (GenBank: GCA_001077635.2) (Perry et al. 2018), and *Deinagkistrodon acutus* (GigaDB: http://dx.doi.org/10.5524/100196) (Yin et al. 2016), 5 coronaviruses with documented origins in bats (bat-SL-CoVZC45 (GenBank: MG772933.1) (Hu et al. 2018), bat-SL-CoVZXC21 (GenBank: MG772934.1) (Hu et al. 2018), BM48-31/BGR/2008 (GenBank: GU190215.1) (Drexler et al. 2010), YNLF_31C (GenBank: KP886808.1), and YNLF_34C (GenBank: KP886809.1)), and 7 additional eukaryotes each known to possess a relatively low genomic GC content (*Lottia gigantea* (GenBank: GCA_000327385.1) (Simakov et al 2013), *Trichoplax adhaerens* (GenBank: GCF_000150275.1) (Srivastava et al. 2008), *Octopus bimaculoides* (GenBank: GCF_001194135.1) (Albertin et al. 2015), *Sarcophilus harrisii* (GenBank: GCF_902635505.1) (Murchison et al. 2012), *Tetranychus urticae* (GenBank: GCF_000239435.1) (Grbić et al. 2011), *Pediculus humanus* (GenBank: GCF_000006295.1) (Kirkness et al. 2010), and *Apis melilifera* (GenBank: GCF_003254395.2) (Wallberg et al. 2019)). For each species, coding sequence GC content was calculated using statswrapper.sh from BBMap v38.75 (Bushnell et al. 2014), and relative synonymous codon usage (RSCU) was calculated using codonw (Peden, 1997), following Ji et al (2020). Output from codonw was parsed using codonw-parser (https://github.com/juliambrosman/parse-codonw) and combined with RSCU values published in Ji et al. A heatmap of RSCU values was generated using pheatmap (Kolde 2015), with species clustered using the default Euclidean method. Additionally, principle components analysis of RSCU values was conducted on the full dataset as well as a subset of focal species (all snake species, one bat, known bat-derived coronaviruses, and the SARS-CoV-2). Squared Euclidian distance (SED) between RSCU values were calculated between SARS-CoV-2 and all other species, and a simple linear regression was used to test for correlation between SED and relative similarity in coding sequence GC content (measured as the difference in coding sequence GC content between SARS-CoV-2 and a given species).

### 4. Accession Numbers of Sequences

| | | | | SAMPLING TIME | | |
|---|---|---|---|---|---|---|
| **Virus_name** | **Species** | **Sample_location** | **Accession_no.** | **YEAR** | **MONTH** | **DAY** |
| RpShaanxi2011 | R_pusillus | Shaanxi | JX993987 | 2011 | 9 | NA |
| HuB2013 | R_sinicus | Hubei | KJ473814 | 2013 | 4 | NA |
| 279_2005 | R_macrotis | Hubei | DQ648857 | 2004 | 11 | NA |
| Rm1 | R_macrotis | Hubei | DQ412043 | 2004 | 11 | NA |
| JL2012 | R_ferrumequinum | Jilin | KJ473811 | 2012 | 10 | NA |
| JTMC15 | R_ferrumequinum | Jilin | KU182964 | 2013 | 10 | NA |
| HeB2013 | R_ferrumequinum | Hebei | KJ473812 | 2013 | 4 | NA |
| SX2013 | R_ferrumequinum | Shanxi | KJ473813 | 2013 | 11 | NA |
| Jiyuan-84 | R_ferrumequinum | Henan-Jiyuan | KY770860 | 2012 | NA | NA |
| Rf1 | R_ferrumequinum | Hubei-Yichang | DQ412042 | 2004 | 11 | NA |
| GX2013 | R_sinicus | Guangxi | KJ473815 | 2012 | 11 | NA |
| Rp3 | R_pearsoni | Guangxi-Nanning | DQ071615 | 2004 | 12 | NA |
| Rf4092 | R_ferrumequinum | Yunnan-Kunming | KY417145 | 2012 | 9 | 18 |
| Rs4231 | R_sinicus | Yunnan-Kunming | KY417146 | 2013 | 4 | 17 |
| WIV16 | R_sinicus | Yunnan-Kunming | KT444582 | 2013 | 7 | 21 |
| Rs4874 | R_sinicus | Yunnan-Kunming | KY417150 | 2013 | 7 | 21 |
| YN2018B | R_affinis | Yunnan | MK211376 | 2016 | 9 | NA |
| Rs7327 | R_sinicus | Yunnan--Kunming | KY417151 | 2014 | 10 | 24 |
| Rs9401 | R_sinicus | Yunnan-Kunming | KY417152 | 2015 | 10 | 16 |
| Rs4084 | R_sinicus | Yunnan-Kunming | KY417144 | 2012 | 9 | 18 |
| RsSHC014 | R_sinicus | Yunnan-Kunming | KC881005 | 2011 | 4 | 17 |
| Rs3367 | R_sinicus | Yunnan-Kunming | KC881006 | 2012 | 3 | 19 |
| WIV1 | R_sinicus | Yunnan-Kunming | KF367457 | 2012 | 9 | NA |
| YN2018C | R_affinis | Yunnan-Kunming | MK211377 | 2016 | 9 | NA |
| As6526 | Aselliscus_stoliczkanus | Yunnan-Kunming | KY417142 | 2014 | 5 | 12 |
| YN2018D | R_affinis | Yunnan | MK211378 | 2016 | 9 | NA |
| Rs4081 | R_sinicus | Yunnan-Kunming | KY417143 | 2012 | 9 | 18 |
| Rs4255 | R_sinicus | Yunnan-Kunming | KY417149 | 2013 | 4 | 17 |
| Rs4237 | R_sinicus | Yunnan-Kunming | KY417147 | 2013 | 4 | 17 |
| Rs4247 | R_sinicus | Yunnan-Kunming | KY417148 | 2013 | 4 | 17 |
| Rs672 | R_sinicus | Guizhou | FJ588686 | 2006 | 9 | NA |
| YN2018A | R_affinis | Yunnan | MK211375 | 2016 | 9 | NA |
| YN2013 | R_sinicus | Yunnan | KJ473816 | 2010 | 12 | NA |
| Anlong-103 | R_sinicus | Guizhou-Anlong | KY770858 | 2013 | NA | NA |
| Anlong-112 | R_sinicus | Guizhou-Anlong | KY770859 | 2013 | NA | NA |
| HSZ-Cc | SARS-CoV-1 | Guangzhou | AY394995 | 2002 | NA | NA |
| YNLF_31C | R_Ferrumequinum | Yunnan-Lufeng | KP886808 | 2013 | 5 | 23 |
| YNLF_34C | R_Ferrumequinum | Yunnan-Lufeng | KP886809 | 2013 | 5 | 23 |
| F46 | R_pusillus | Yunnan | KU973692 | 2012 | NA | NA |
| SC2018 | R_spp | Sichuan | MK211374 | 2016 | 10 | NA |
| LYRa11 | R_affinis | Yunnan-Baoshan | KF569996 | 2011 | NA | NA |
| Yunnan2011 | Chaerephon_plicata | Yunnan | JX993988 | 2011 | 11 | NA |
| Longquan_140 | R_monoceros | China | KF294457 | 2012 | NA | NA |
| HKU3-1 | R_sinicus | Hong_Kong | DQ022305 | 2005 | 2 | 17 |
| HKU3-3 | R_sinicus | Hong_Kong | DQ084200 | 2005 | 3 | 17 |
| HKU3-2 | R_sinicus | Hong_Kong | DQ084199 | 2005 | 2 | 24 |
| HKU3-4 | R_sinicus | Hong_Kong | GQ153539 | 2005 | 7 | 20 |
| HKU3-5 | R_sinicus | Hong_Kong | GQ153540 | 2005 | 9 | 20 |
| HKU3-6 | R_sinicus | Hong_Kong | GQ153541 | 2005 | 12 | 16 |
| HKU3-10 | R_sinicus | Hong_Kong | GQ153545 | 2006 | 10 | 28 |
| HKU3-9 | R_sinicus | Hong_Kong | GQ153544 | 2006 | 10 | 28 |

| | | | | | | |
|---|---|---|---|---|---|---|
| HKU3-11 | R_sinicus | Hong_King | GQ153546 | 2007 | 3 | 7 |
| HKU3-13 | R_sinicus | Hong_Kong | GQ153548 | 2007 | 11 | 15 |
| HKU3-12 | R_sinicus | Hong_Kong | GQ153547 | 2007 | 5 | 15 |
| HKU3-7 | R_sinicus | Guangdong | GQ153542 | 2006 | 2 | 15 |
| HKU3-8 | R_sinicus | Guangdong | GQ153543 | 2006 | 2 | 15 |
| CoVZC45 | R_sinicus | Zhoushan-Dinghai | MG772933 | 2017 | 2 | NA |
| CoVZXC21 | R_sinicus | Zhoushan-Dinghai | MG772934 | 2015 | 7 | NA |
| Wuhan-Hu-1 | SARS-CoV-2 | Wuhan | MN908947 | 2019 | 12 | NA |
| BtKY72 | R_spp | Kenya | KY352407 | 2007 | 10 | NA |
| BM48-31 | R_blasii | Bulgaria | NC_014470 | 2008 | 4 | NA |
| RaTG13 | R_affinis | Yunnan | EPI_ISL_402131 | 2013 | 7 | 24 |
| P4L | pangolin | Guangxi | EPI_ISL_410538 | 2017 | NA | NA |
| P5L | pangolin | Guangxi | EPI_ISL_410540 | 2017 | NA | NA |
| P5E | pangolin | Guangxi | EPI_ISL_410541 | 2017 | NA | NA |
| P1E | pangolin | Guangxi | EPI_ISL_410539 | 2017 | NA | NA |
| P2V | pangolin | Guangxi | EPI_ISL_410542 | 2017 | NA | NA |
| Pangolin-CoV | pangolin | Guandong | EPI_ISL_410721 | 2019 | 3 | NA |

### 5. Phylogenetic incongruence among 10 breakpoint-free regions (BFRs) built for NRR1

Shown after the references as Figures S5 to S14.

Figure S5
Breakpoint-free region A: nucleotides 13291-19628

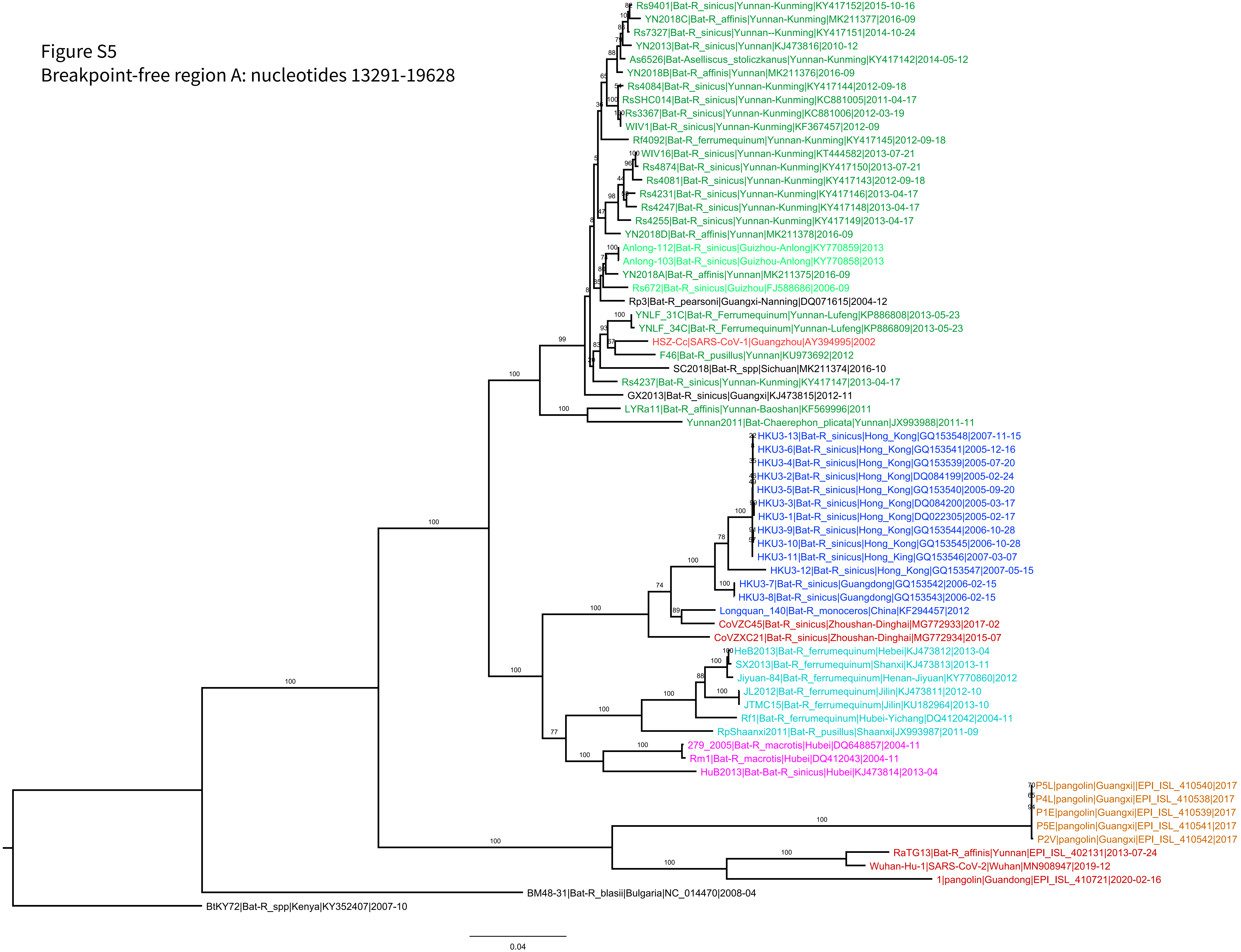

Figure S6
BFR B: 3625-9150

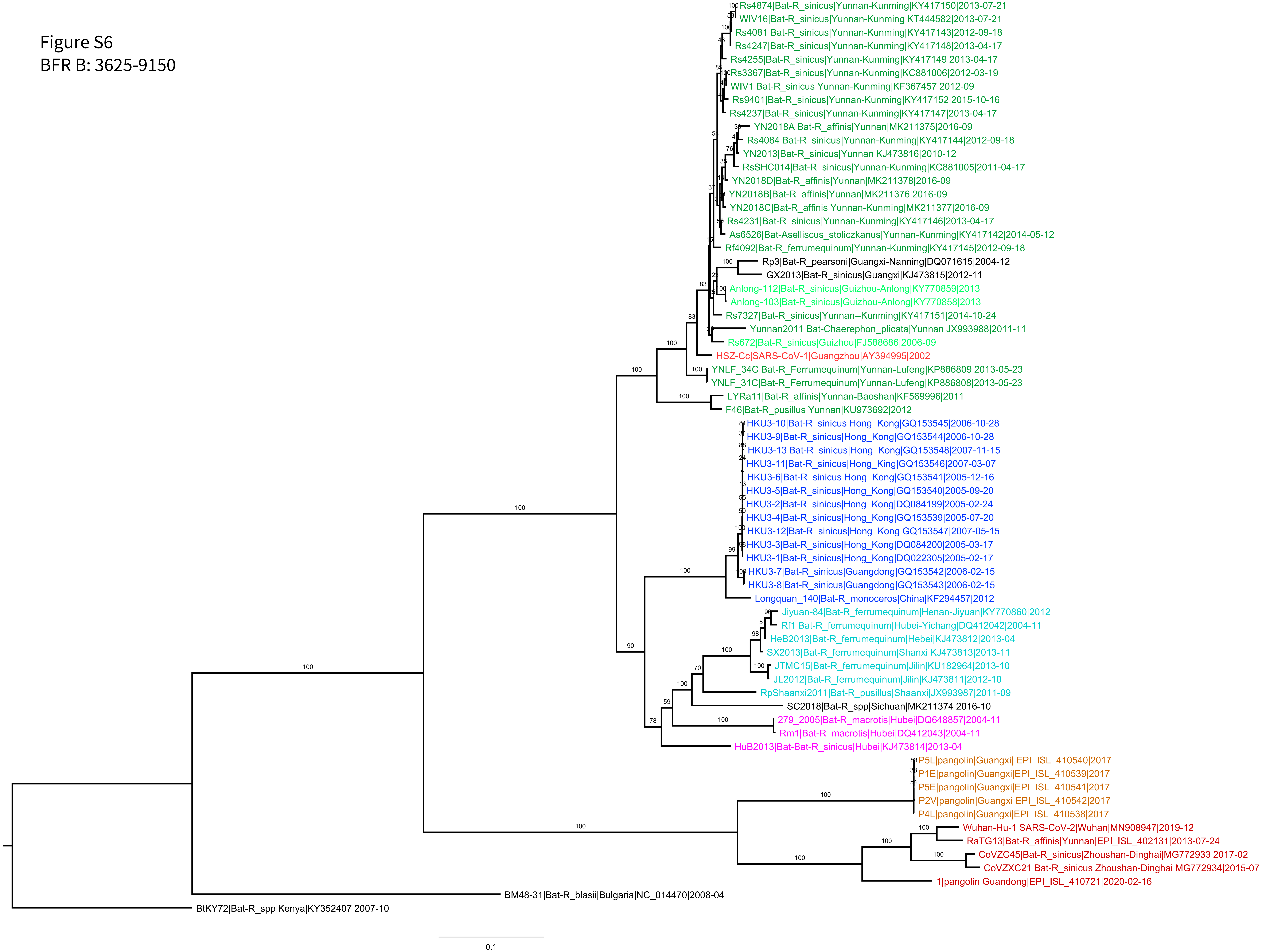

Figure S7
BFR C: 9261-11795

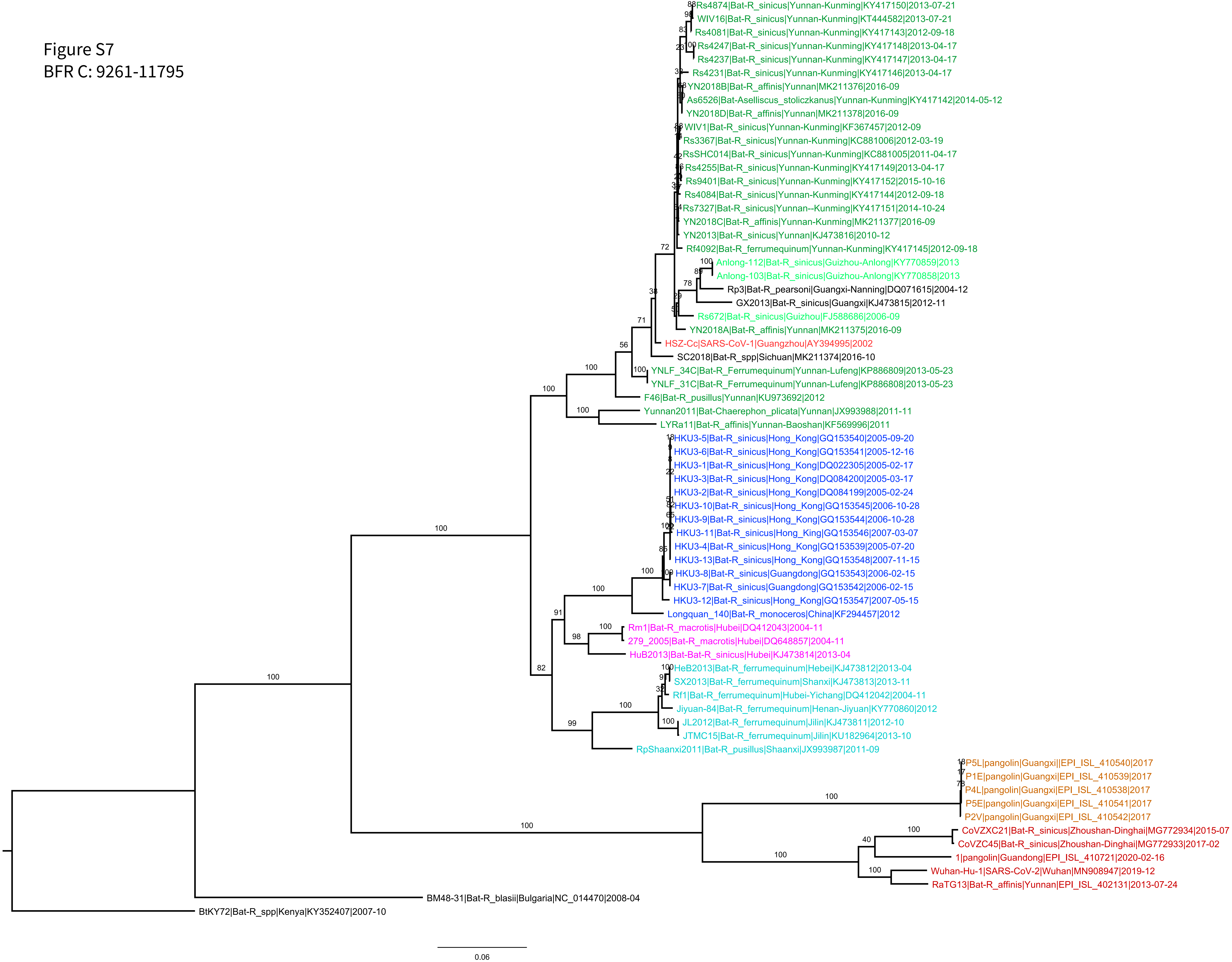

Figure S8

BFR D: 27702-28843

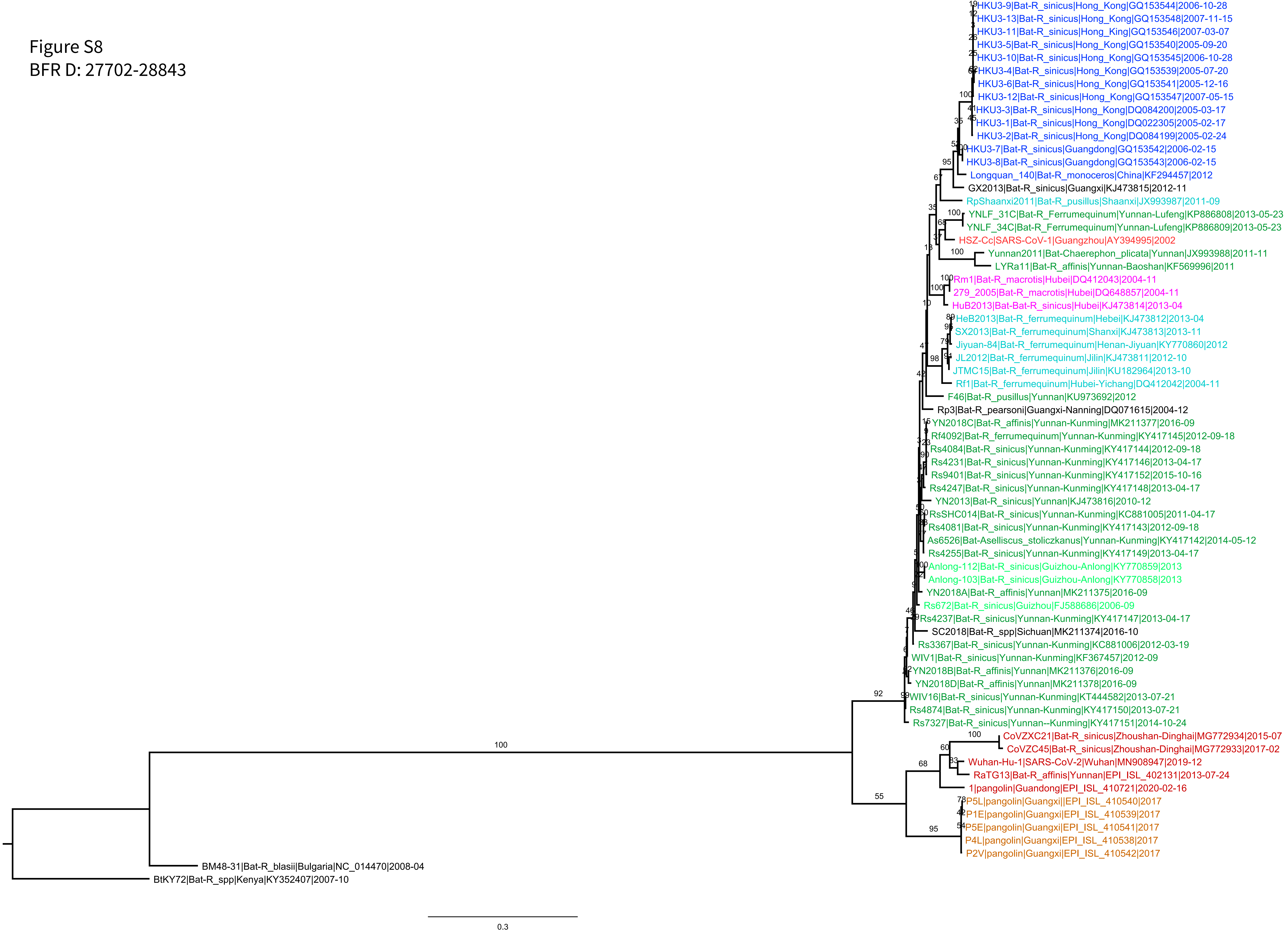

Figure S9
BFR E: 29574-30650

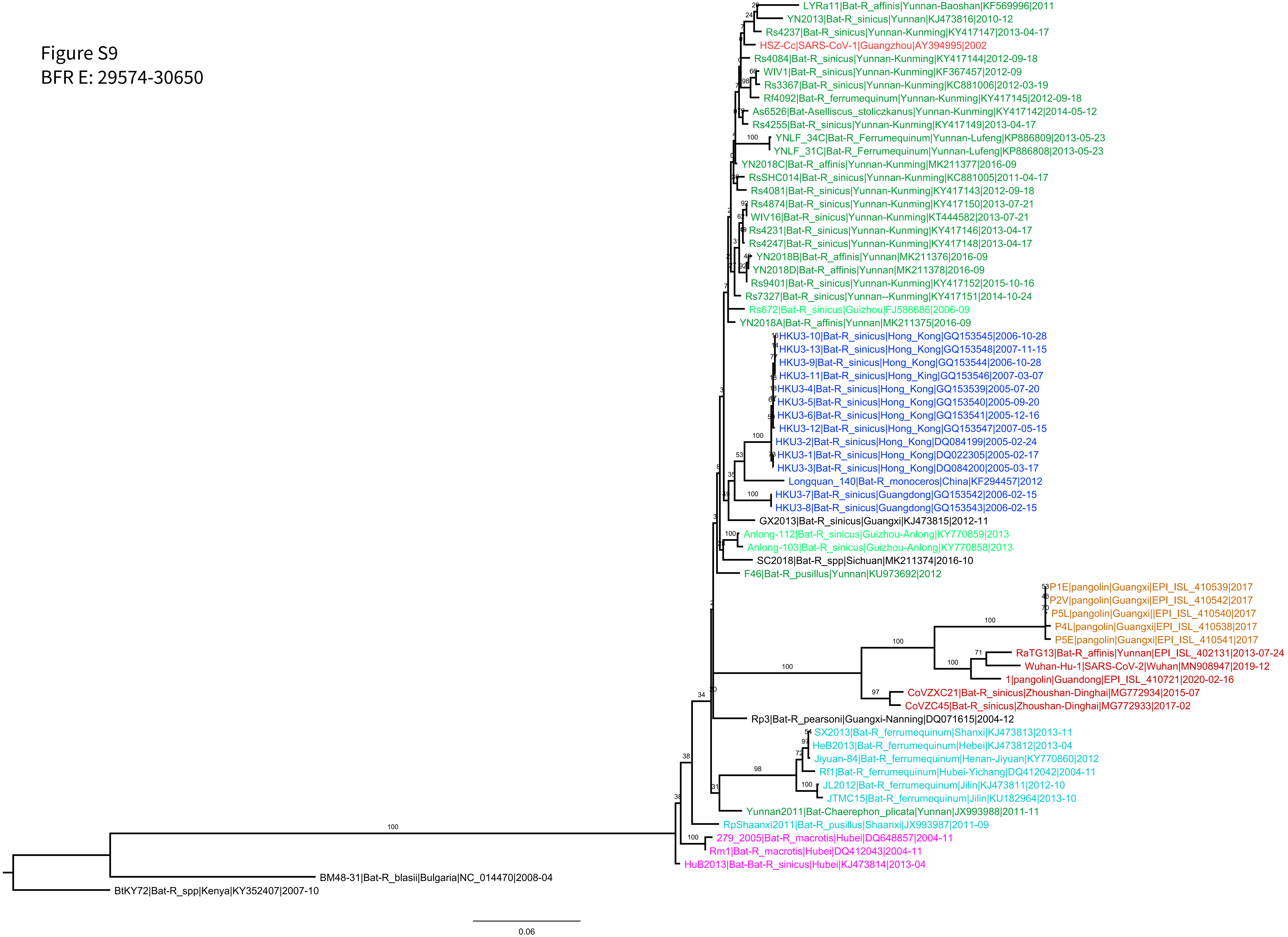

Figure S10
BFR F: 24795-25837

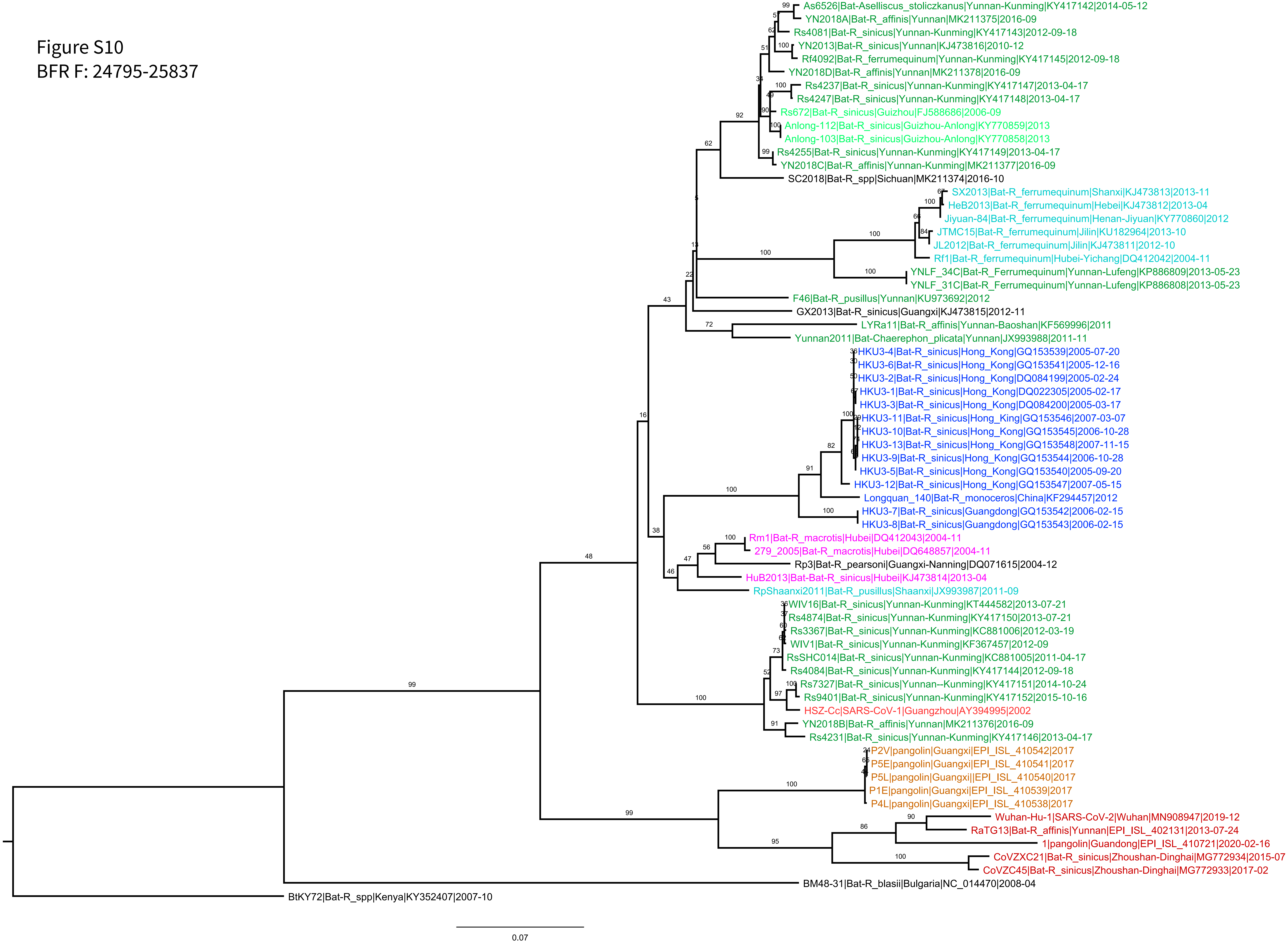

Figure S11
BFR G: 23631-24633

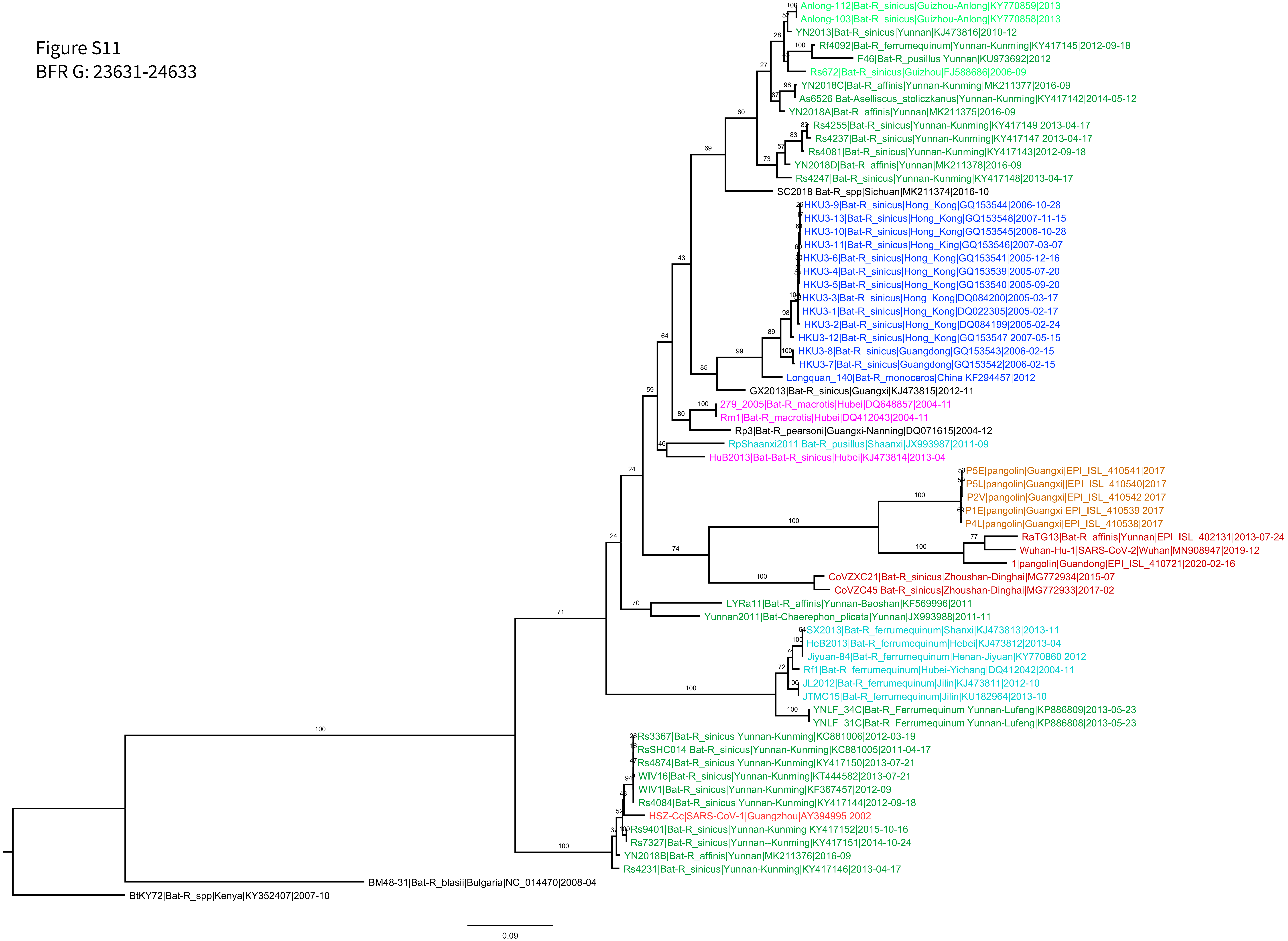

Figure S12
BFR H: 12443-13291

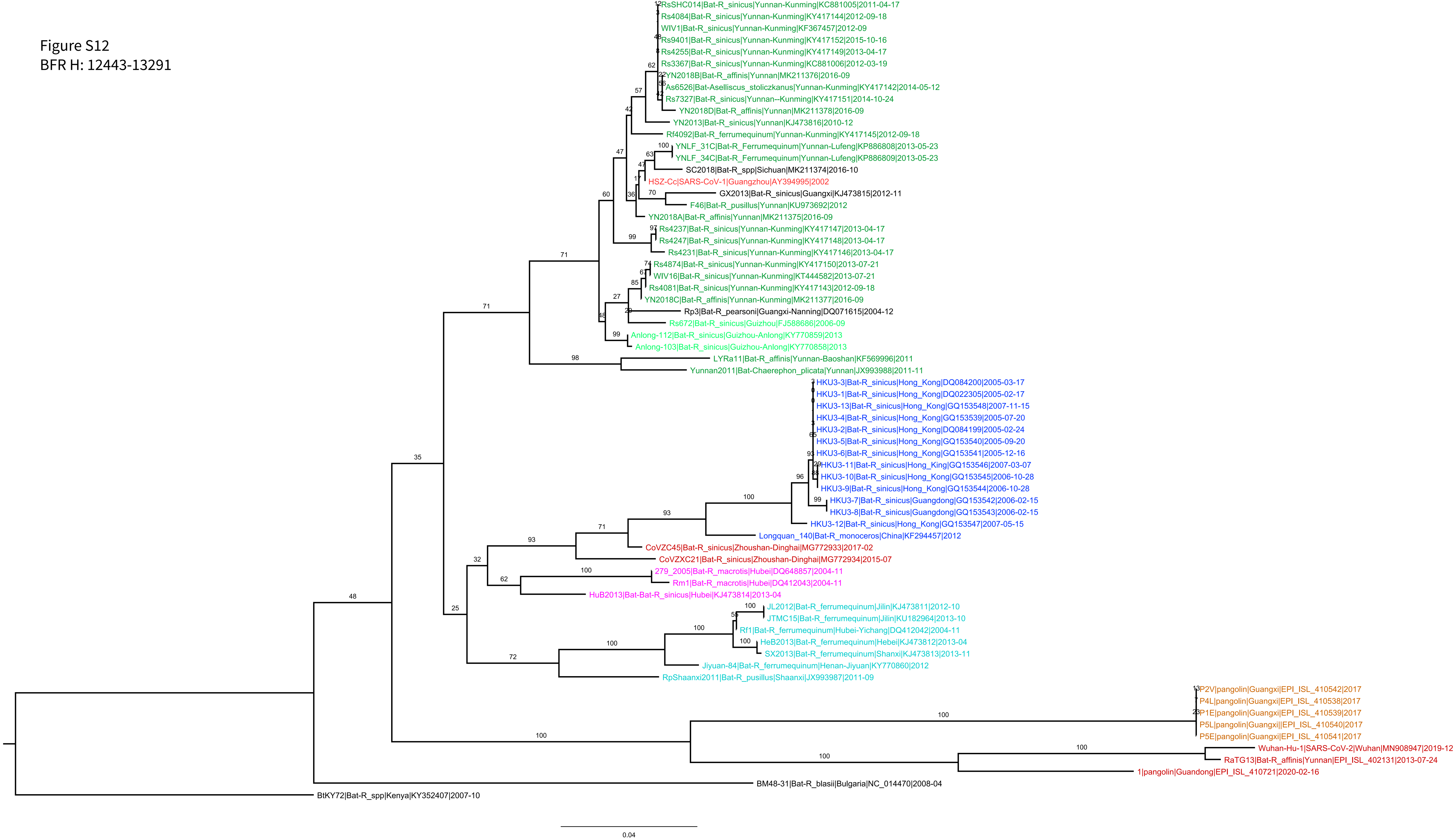

Figure S13
BFR I: 962-1686

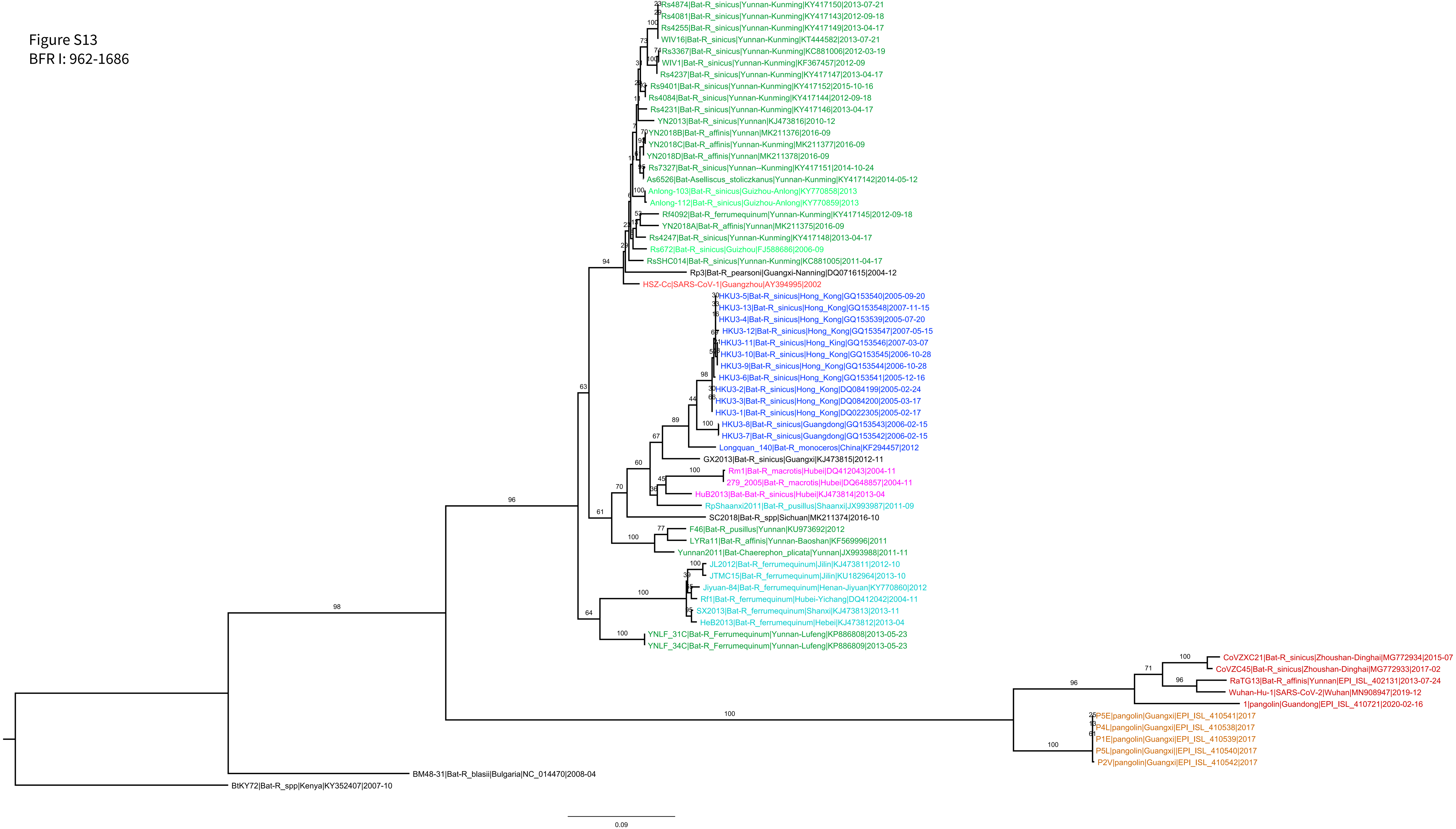

Figure S14
BFR J: 147-695

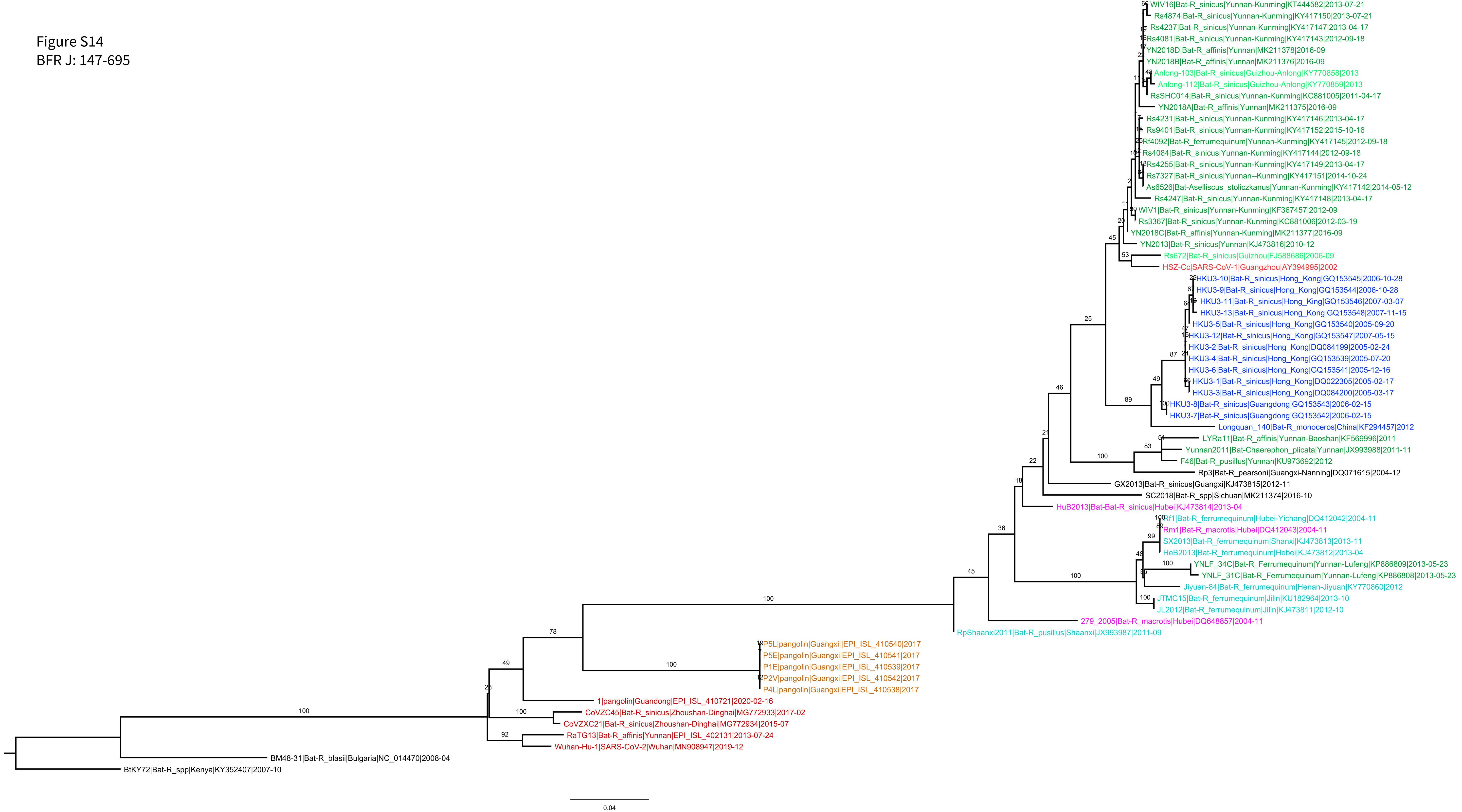
